# Oxygen gradients reshape cross-feeding through emergent spatial organization of gut commensal bacteria

**DOI:** 10.64898/2026.05.02.721930

**Authors:** David Scheidweiler, Elise Bornet, Hannah Ochner, Sonja Blasche, Stanislas Thiriet-Rupert, Louis Dorison, Christophe Beloin, Yohan Davit, Samy Gobaa, Tanmay A. M. Bharat, Alexander J. Westermann, Kiran R. Patil, Jean-Marc Ghigo

## Abstract

Microbial interactions unfold within environments structured by physical transport and chemical gradients. Yet most mechanistic studies rely on well-mixed systems that mask the reciprocal influences of environmental heterogeneity on metabolism and ecology. Here, we investigate how the physical environment modulates the interaction between the gut commensal *Bacteroides thetaiotaomicron* and *Escherichia coli*. In anoxic liquid culture, cell-resolved isotope imaging and genetic perturbations reveal exploitative cross-feeding, where *E. coli* consumes diffusible sugars released by *B*. thetaiotaomicron during starch degradation. When exposed to intestinal-like oxygen gradients in microfluidics, the interaction is restructured by spatial organization. The species self-organize into complementary niches: *E. coli* locally depletes sugars and oxygen, thereby expanding the anoxic niche required by *B. thetaiotaomicron*. A reactive transport model confirms that this organization arises from coupled feedback between physical transport and metabolic reaction rates. Together, our results reveal how physical structure and chemical gradients convert an exploitative cross-feeding interaction into a dynamic niche-construction process that generates emergent spatial organization and stabilizes coexistence.

## Introduction

Microbial communities underpin essential ecosystem functions across natural and host-associated environments, from global biogeochemical cycles to sustaining host health [1–3]. These functions emerge from complex networks of microbial interactions mediated by the surrounding chemical landscape of resources and metabolites [4–8]. Yet this landscape is not a static template but is actively shaped by the interplay between physical structure, transport and microbial metabolism that reconfigures the local environment [9–11]. While such feedbacks are increasingly recognized as key mediators of microbial interactions [12–14], mechanistic insights remain largely confined to well-mixed laboratory systems. By homogenizing the environment, such systems decouple metabolism from its spatial context, leaving the role of physical structure in shaping interaction outcomes underexplored.

Host-associated microbiomes represent a compelling example of spatially structured ecosystems [15, 16]. In the mammalian gut, microbial interactions are constrained by several host-controlled factors, including immune activity, dietary intake, and transit time [17–20]. Dietary and host-derived glycans are primary resources for which microorganisms compete; these complex carbohydrates are degraded by specialized taxa [21, 22], ultimately releasing sugars and metabolites that couple species through metabolic networks [23, 24]. The availability of these metabolites is further modulated by intestinal transit time [25–27], and by steep oxygen gradients arising from the diffusion of oxygen from host tissues [28–30], both of which contribute to the marked microbial compositional differences observed along the intestinal tract [31–33]. Beyond these physical drivers, microbial activity also modifies the local physicochemical condition—for example through oxygen depletion—highlighting a reciprocal feedback between microbial metabolism and the chemical environment, the mechanistic basis of which remains poorly resolved.

Here, we investigate how transport and metabolic-environmental feedbacks govern the interaction between the primary polysaccharide degrader *Bacteroides thetaiotaomicron* and the facultative anaerobe *Escherichia coli*. By integrating microfluidic control of environmental gradients with isotope imaging, transcriptomics, and modeling, we show that the interaction outcomes are a predictable function of the physical environment. We demonstrate that the interplay between flow-mediated transport and metabolic oxygen consumption reconfigures interaction outcomes, shifting the relationship between these species from competitive exploitation toward facilitated coexistence. Together, our results provide a mechanistic framework linking metabolic feedback and environmental structure to the spatial organization of microbial systems in structured habitats.

## Results

### Starch-mediated interactions provide resource access for *E. coli* in liquid co-culture with *B. thetaiotaomicron*

To investigate how the strict anaerobe *B. thetaiotaomicron* and the facultative anaerobe *E. coli* interact in presence of a complex dietary polysaccharide. We grew mono- and co-culture of *B. thetaiotaomicron* VPI-5482 and *E. coli* MG1655 strains in anaerobic condition in defined medium containing, as the sole source of carbon, amylopectin (Fig. 1A) a highly branched glucose polymer entering the colon in the composition of starch. In monoculture, *B. thetaiotaomicron* degraded amylopectin thanks to its glycolytic repertoire and reached a cell density of ~ 10^9^ colony forming units (CFU) mL^−1^, whereas *E. coli*, which cannot hydrolyze amylopectin, did not grow (Fig. 1B). In contrast, in co-culture, *E. coli* benefited from *B. thetaiotaomicron* presence and increased in abundance, while *B. thetaiotaomicron* reached a lower final density as in mono-culture (~ 10^8^ CFU mL^−1^).

**Figure 1:**
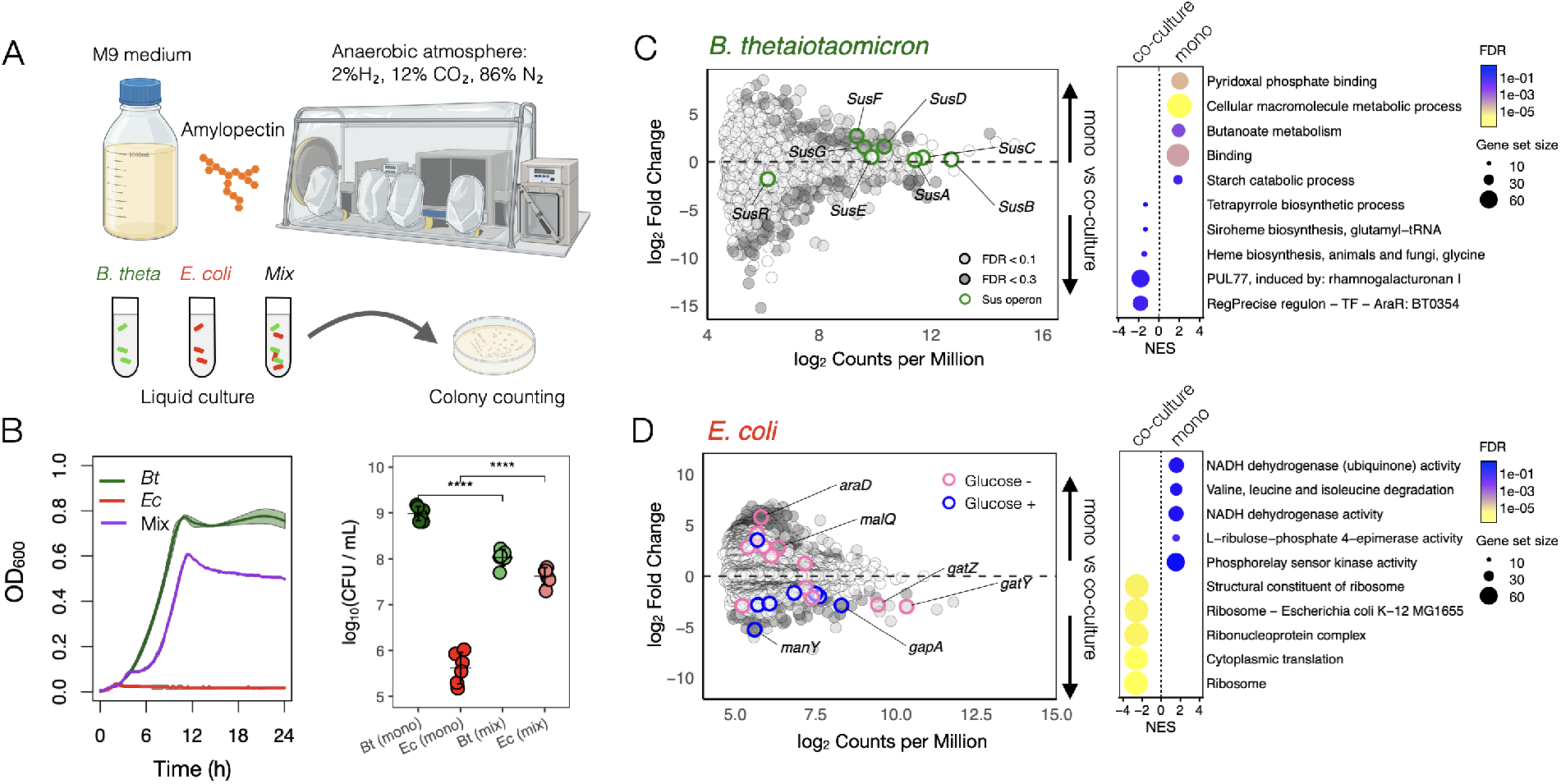
Polysaccharide-mediated interactions between *B. thetaiotaomicron* VPI-5482 and *E. coli* MG1655. (A) Experimental setup. Monocultures of *B. thetaiotaomicron, E. coli*, or a 1:1 co-culture were grown anaerobically in defined M9 medium supplemented with amylopectin as sole carbon source. (B) Left. Growth curves of monocultures and co-culture in amylopectin medium after 24 hours. Lines represent averages of independent experiments (*n* = 3); shaded areas represent standard deviation. Right. Colony-forming units (CFU mL^−1^) showing loss of *E. coli* growth in monoculture and its rescue in co-culture (*n* = 6). Each point represents a single replicate sample, colored by treatment. Significance was determined by two-sided t test. (C) Mean-difference (MD) plot of *B. thetaiotaomicron* transcriptomes comparing monoculture (top) and co-culture (bottom) after 24 hours of growth (*n* = 3). The x-axis shows the average expression level of each gene as log_2_ counts per million, and the y-axis shows the log_2_ fold change in expression between monoculture and co-culture. Genes with a false discovery rate (FDR) *<* 0.1 are highlighted in light gray, and those with FDR *<* 0.3 in dark gray; Sus-operon and Sus-homolog genes are indicated. (right) Enrichment analysis of *B. thetaiotaomicron* genes in mono- and co-culture. Dot size reflects gene set size, and color indicates normalized enrichment score (NES). (D) MD plot of *E. coli* transcriptomes comparing monoculture (top) and co-culture (bottom) after 24 hours of growth (*n* = 3). Genes induced in the presence of glucose are shown in blue, while genes repressed by glucose are shown in pink. (right) Enrichment analysis of *E. coli* genes in mono- and co-culture. Dot size reflects gene set size, and color indicates normalized enrichment score (NES).

To better understand how this interaction unfolds at the molecular level, we examined changes in the two species’ transcriptional response following a 24 hour incubation. Transcriptomic profiling of *B. thetaiotaomicron* revealed coordinated changes within the genes encoding the amylopectin-degrading Sus locus BT3698-3705 (PUL66), one of the many *B. thetaiotaomicron* Polysaccharide Utilization Loci (PUL) glycan-uptake systems (Fig. 1C). When compared to co-culture, monocultured *B. thetaiotaomicron* upregulates the genes needed to bind amylopectin at the cell surface—primarily *susD* (*BT3701*)—and to initiate its extracellular degradation through the outer-membrane hydrolase SusG (BT3698). This activity releases glucose and malto-oligosaccharides into the surrounding medium prior to import through the TonB-dependent transporter SusC (BT3702) [34]. In contrast, the divergently transcribed regulatory gene *susR* (*BT3705*), whose expression responds to periplasmic starch-derived oligosaccharides, was more expressed in co-culture (−1.8 log_2_ fold change), suggesting that the oligosaccharide environment sensed by *B. thetaiotaomicron* is altered in the presence of *E. coli*. Beyond the core Sus operon, additional Sus homologs were induced in co-culture, indicating a broader reorganization of polysaccharide catabolism (SI, Fig. S6A); consistent with this, gene set enrichment analysis revealed coordinated activation of starch catabolic and associated biosynthetic pathways, pointing to a global metabolic adjustment to altered substrate availability (Fig. 1C).

While several *E. coli* genes did not meet stringent false discovery rate thresholds (FDR ≤0.2–0.3) due to low-input RNA-seq, such data remain informative for hypothesis generation and pathway-level analyses [35]. Consistent with this, genes associated with glucose uptake and metabolism were upregulated in *E. coli* during co-culture, including the sugar-specific phosphotransferase component gene *manY* and the glycolytic enzyme gene *gapA* (Fig. 1D, blue). In parallel, operons typically induced under glucose-limiting conditions—such as *araD* and maltodextrin-associated genes (*malQ*)—were repressed (Fig. 1D, pink), consistent with glucose-dependent carbon catabolite regulation. At the pathway level, gene set enrichment analysis supported this interpretation, revealing enrichment of translation, central metabolism, and respiratory functions, indicative of a growth-optimized physiological state in co-culture (Fig. 1D). Taken together, these data indicate that *E. coli* preferentially exploits starch-derived sugars released by *B. thetaiotaomicron*, reinforcing carbon catabolite repression and suppressing the utilization of alternative carbon sources.

### *E. coli* uses maltose and glucose transport to cross-feed on public good metabolites released by *B. thetaiotaomicron*

To determine whether *E. coli* benefits from enhanced access to pre-existing nutrients in the defined medium or rather from newly accumulating metabolites released during *B. thetaiotaomicron*-mediated polysaccharide breakdown, we cultured the bacteria in M9 medium supplemented with ^13^C-labeled starch. Combining fluorescence microscopy, cryo-EM and chemical imaging using focused ion beam secondary ion mass spectrometry (cryo-CLEM-FIB-SIMS) [37], we observed a clear isotopic enrichment in *E. coli* cells only in co-culture with *B. thetaiotaomicron* (Fig. 2A, SI Fig. S1). This provides direct evidence that *E. coli* incorporates ^13^C originating from *B. thetaiotaomicron*-mediated starch degradation, demonstrating a trophic interaction in which *E. coli* growth is fueled by polysaccharide-derived metabolites.

**Figure 2:**
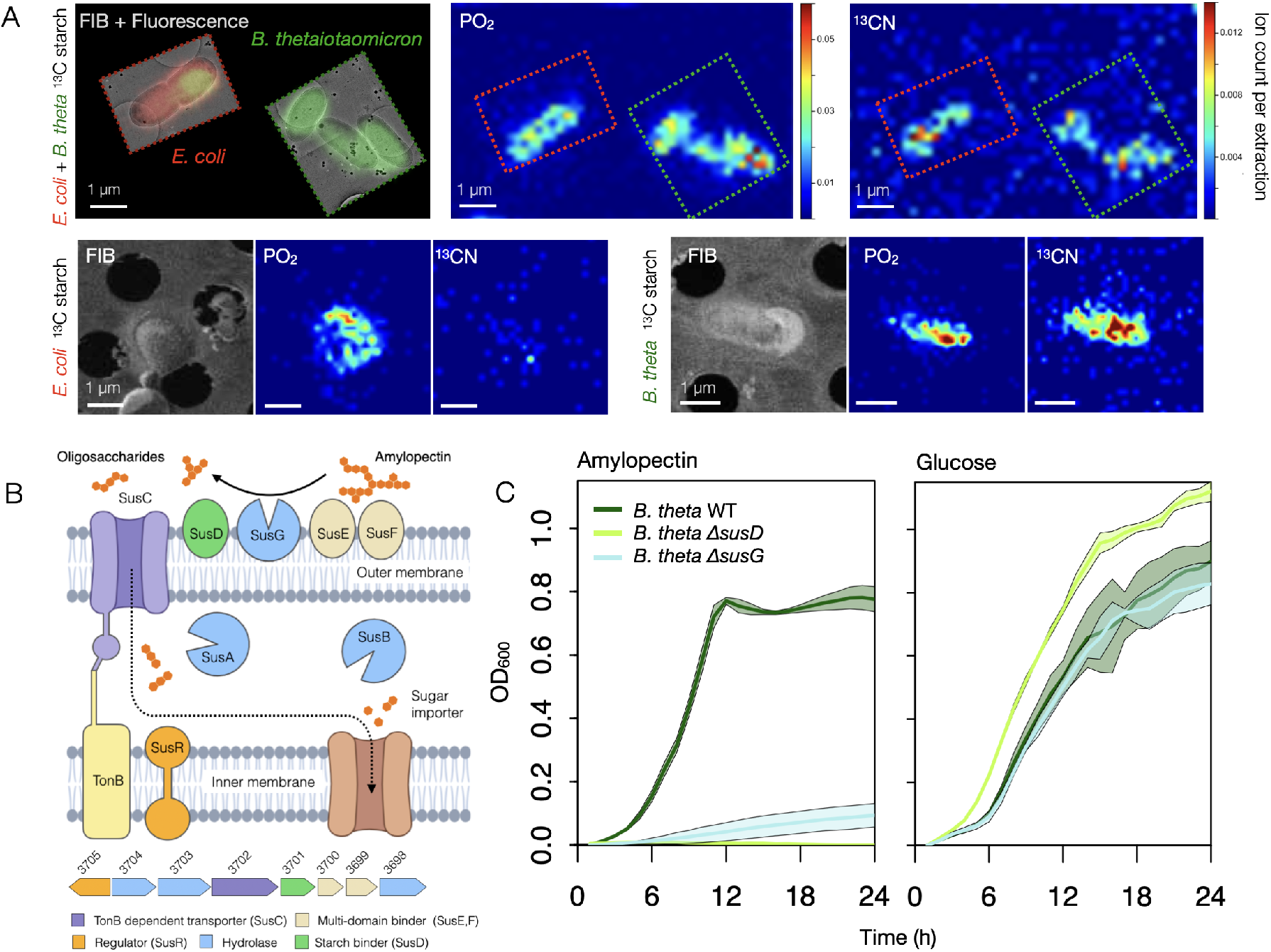
Chemical mapping reveals a Sus-dependent trophic interaction. (A) Cryo-CLEM-FIB-SIMS analysis of a co-culture containing both *B. thetaiotaomicron* and *E. coli*, grown in medium supplemented with ^13^C-labelled starch. Due to fluorescent labelling, the two species are distinguishable in cryo-LM (*B. thetaiotaomicron* green, *E. coli* red, top left). The corresponding cryo-EM images are overlaid with the fluorescent signal. Cryo-FIB-SIMS analysis shows ^13^CN signal from both *B. thetaiotaomicron* and *E. coli* cells. A strong PO_2_ SIMS signal is observed in both cell types (a signal found in all cells [36]), shown here to clarify the location of the cells in the SIMS data. For comparison, SIMS data of monocultures of *B. thetaiotaomicron* and *E. coli* grown in ^13^C-labelled starch, respectively, is shown in the bottom panels. The secondary electron FIB images (left panels) are additionally shown to visualize the location of the cells. While cells of both cultures show clear PO_2_ signal, only *B. thetaiotaomicron* monoculture presents a strong ^13^CN signal, whereas *E. coli* does not. SIMS images for each ionic species have been scaled to the same color scale, representing ion count per extraction, to allow comparison. (B) Top, schematic of the *B. thetaiotaomicron* starch utilization system (Sus), showing extracellular binding, hydrolysis, and transport of malto-oligosaccharide, adapted from [34]. Bottom, full amylopectin-degrading Sus locus BT3698-3705 (PUL66). (C) Growth of *B. thetaiotaomicron* wild type, Δ*susD* (BT3701), and Δ*susG* (BT3698) mutants on amylopectin (left) and glucose (right). Lines represent averages of independent experiments (*n* = 3); shaded areas represent standard deviation.

As *E. coli* benefits from *B. thetaiotaomicron* starch hydrolysis, we next verified that *B. thetaiotaomicron*’s canonical starch-utilization machinery is required under our experimental conditions (Fig. 2B). We showed that the growth of isogenic *B. thetaiotaomicron* Δ*susD* and Δ*susG* mutants were indeed markedly impaired on amylopectin while growing normally on glucose (Fig. 2C), confirming that SusD-mediated polysaccharide capture and SusG-dependent extracellular cleavage are essential for amylopectin catabolism in M9 medium.

We then tested whether *E. coli* relies on starch-derived breakdown products by disrupting its major glucose and maltose utilization pathways. For this we deleted the genes encoding the glucose-specific phosphotransferase system (Δ*ptsI* Δ*ptsH* Δ*crr*, hereafter Δ*pts*), the maltose-binding protein required for malto-oligosaccharide import (Δ*malE*), or all these genes (Δ*pts* Δ*malE*) (Fig. 3A). These strains displayed impaired growth in the defined medium amended with glucose, maltose and amylopectin (Fig. 3B), but retained normal growth in LB, an amino-acid–rich medium (SI, Fig. S2). We next generated *B. thetaiotaomicron* spent medium using amylopectin as the sole carbon source and eventually equilibrated it to either oxic or anoxic conditions (Fig. 3C). When grown in anoxic spent medium, *E. coli* reached a lower cell density than under oxic conditions, and only the wild type proliferated efficiently, while the Δ*pts* Δ*malE* mutant showed no growth and the Δ*malE* displayed a strong growth defect (Fig. 3D). In the anoxic condition, the Δ*pts* mutant with a slower growth rate ultimately reached a similar density as the wild type after ~ 12 hours (Fig. 3D). Taken together, these results provide strong evidence that *E. coli* relies on maltose/glucose transport pathways to exploit the metabolites released by *B. thetaiotaomicron*, and that oxygen availability modulates the energetic payoff of this cross-feeding interaction.

**Figure 3:**
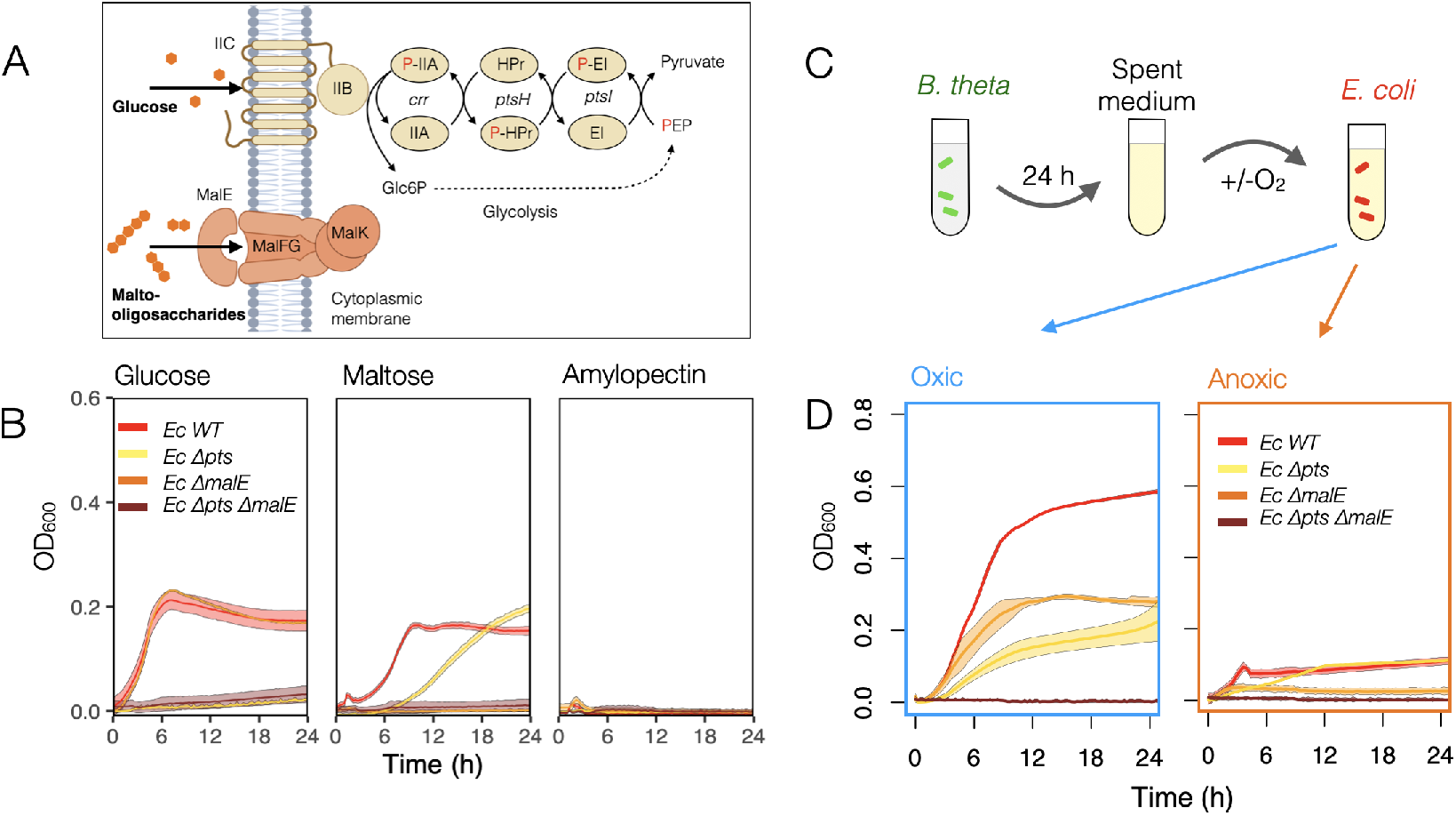
Glucose and maltose uptake pathways determine *E. coli* growth on polysaccharide-derived metabolites. (A) Schematic of *E. coli* carbohydrate uptake pathways, showing the glucose phosphotransferase system (PTS) and the maltose/maltodextrin transporter complex MalEFGK. Adapted from [38, 39]. (B) Growth of *E. coli* wild type, Δ*pts*, Δ*malE*, and Δ*pts*Δ*malE* mutants in anoxic M9 medium supplemented with glucose, maltose, or amylopectin as the carbon source. Lines represent averages of independent experiments (*n* = 3); shaded areas represent standard deviation. (C) Experimental setup for spent medium assays. *B. thetaiotaomicron* was grown for 24 h in amylopectin medium; cell-free spent medium was then used to culture *E. coli* strains under oxic or anoxic conditions. (D) Growth of *E. coli* wild type and carbohydrate-uptake deficient mutants in *B. thetaiotaomicron*-spent medium under oxic (left) or anoxic (right) conditions. Lines represent averages of independent experiments (*n* = 3); shaded areas represent standard deviation.

### Oxygen gradients promote spatial organization of *B. thetaiotaomicron* and *E. coli*

While the described liquid culture experiments established that *B. thetaiotaomicron* releases diffusible metabolites supporting *E. coli* growth, this interaction took place in a well-mixed environment without any chemical gradients. However, microorganisms inhabit environments characterized by spatial constraints and physicochemical gradients that can profoundly alter metabolic interactions. In the mammalian intestine for instance, oxygen exhibits extremely steep gradients: concentrations fluctuate near the mucosal surface, due to epithelial metabolism and inflammation, yet drop to complete anoxia only a few hundred micrometers into the lumen [40]. We therefore investigated whether the cross-feeding observed in bulk liquid co-culture persists when *B. thetaiotaomicron* and *E. coli* interact in a spatially structured microenvironment that imposes flow and oxygen gradients.

We engineered a microfluidic device featuring a central lumen channel flanked by crypt-like side cavities, in which the oxygen-depleted M9 medium supplemented with amylopectin was perfused through the central channel (Fig. 4A, *Q* = 0.5 *µ*L min^−1^, *v* = 50 *µ*m s^−1^). The oxygen gradient along the orthogonal crypt-like cavities was controlled using a microscope-stage CO_2_–O_2_ incubator, allowing diffusion of gas through the oxygen-permeable PDMS membrane and reproducing physiological oxygen concentrations near the intestinal mucosa [40]. Due to the lumen–crypt structure, the competition between advective and diffusive transport time scales leads to a steep 0–0.5% oxygen gradient along the crypt axis, as confirmed by COMSOL simulations (Fig. 4A).

**Figure 4:**
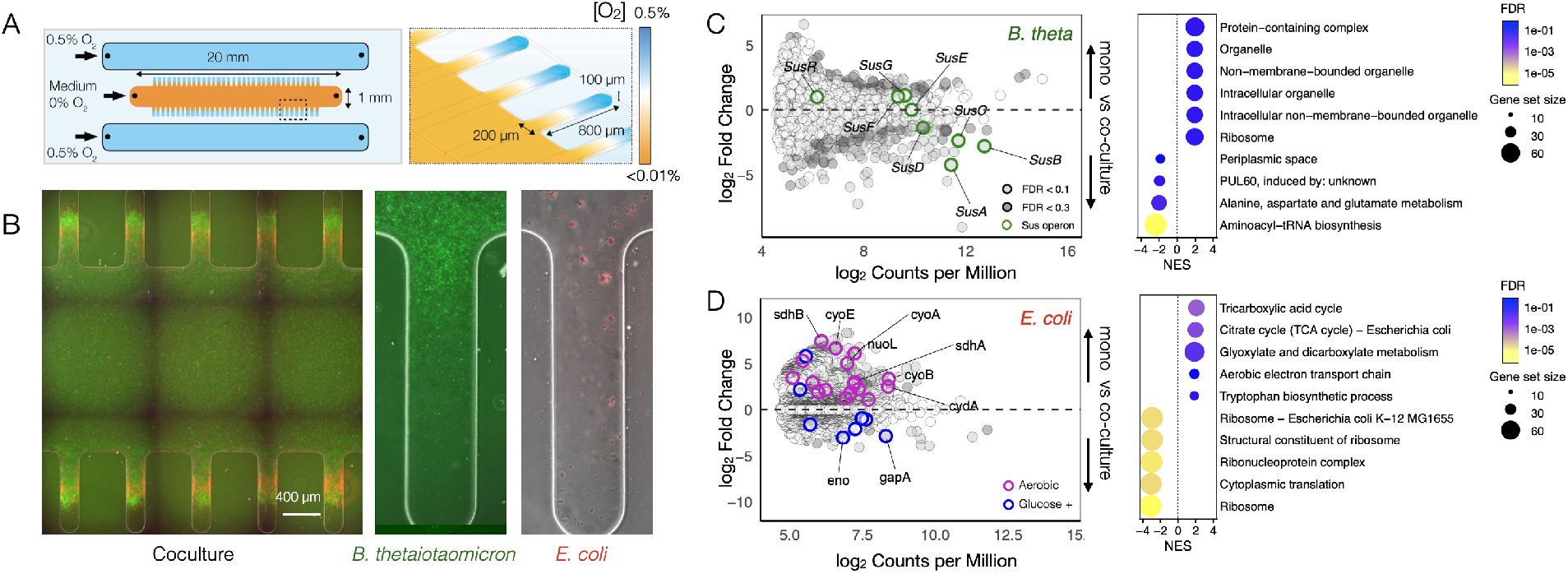
Imposing oxygen gradient in microfluidic devices. (A) Left. Schematic of the microfluidic device used to generate a controlled oxygen gradient along crypt-like structures. Oxygen-depleted M9 medium supplemented with amylopectin was perfused through the central channel (*v* ≈ 50 *µ*m s^−1^), while lateral channels exposed to defined oxygen concentrations established a diffusive gradient across the cavities. Right, numerical simulations illustrate the predicted steady-state oxygen distribution in the chip. (B) Fluorescence images overlaid on transmitted-light micrographs show the spatial organization of *B. thetaiotaomicron* (green) and *E. coli* (red) in co-culture or monoculture after 24 h of growth in the microfluidic device. Scale bar = 400 *µ*m. (C) Mean-difference (MD) plot of *B. thetaiotaomicron* transcriptomes comparing monoculture (top) and co-culture (bottom) after 24 h in the microfluidic device (*n* = 3). The x-axis shows the average expression level of each gene as log_2_ counts per million, and the y-axis shows the log_2_ fold change in expression between monoculture and co-culture. Genes with false discovery rate (FDR) *<* 0.1 are shown in light gray, and genes with FDR *<* 0.3 in dark gray; *sus*-operon genes are indicated in green. (right) Enrichment analysis of *B. thetaiotaomicron* genes in mono- and co-culture. Dot size reflects gene set size, and color indicates normalized enrichment score (NES). (D) MD plot of *E. coli* transcriptomes under the same conditions (*n* = 3). Genes involved in glucose utilization are highlighted in blue, and genes encoding components of the aerobic respiratory chain are shown in purple. (right) Enrichment analysis of *E. coli* genes in mono- and co-culture. Dot size reflects gene set size, and color indicates normalized enrichment score (NES).

After 24 h of experiment, the two strains exhibited distinct spatial organization, as visualized by fluorescence microscopy. In monocultures, *B. thetaiotaomicron* colonized the lumen and the entrances of the crypts, while *E. coli* did not grow (Fig. 4B, SI Fig. S3). In co-culture, neither species followed its monoculture pattern: *E. coli* expanded through the crypts, while *B. thetaiotaomicron* penetrated deeper into crypt regions, and reached a higher CFU than *E. coli*. This contrasts the result obtained in oxic liquid culture where *E. coli* outcompetes *B. thetaiotaomicron* (SI, Fig. S4 and S7). The spatial organization was presumably enabled by the accumulation of starch-derived sugars in the crypts, released by *B. thetaiotaomicron*, and by oxygen consumption by *E. coli*, which extended the range of anoxic niches accessible to *B. thetaiotaomicron*. Consistent with this interpretation, in presence of uniformly supplied glucose, both species exhibited a uniform spatial colonization in co-culture (SI, Fig. S5), indicating that in presence of amylopectin, the organization observed depends on localized sugar release rather than oxygen gradients alone.

### Transcriptional reprogramming emerges under spatial structure

To contextualize these spatial phenotypes at the transcriptional level, we recovered the biomass reached in parallel experiments after 24 h from the microfluidic device for ultrasensitive RNA-seq analysis [35]. In co-culture, *B. thetaiotaomicron* showed strong upregulation of genes encoding core components of the amylopectin-degrading Sus system, including *susD* (*BT3701*), *susE* (*BT3702*), *susF* (*BT3703*), and that of the outer-membrane hydrolase BT3704 (Fig. 4C). Notably, several tRNAs were expressed at higher levels in co-culture than in monoculture, a pattern often associated with elevated translational activity and faster growth (SI, Fig. S6A). In turn, *E. coli* showed an increased expression of glucose-supported central metabolism in co-culture, including glycolytic genes such as *gapA* and *eno* (Fig. 4D, blue). Conversely, multiple oxygen-dependent respiratory modules were reduced in co-culture relative to monoculture (Fig. 4D, purple), including components of the aerobic electron transport chain (e.g., *cyoA, cyoB, cyoE* and *nuoL*) and the TCA-linked dehydrogenases *sdhA* and *sdhB*. Together, this coordinated shift is consistent with *E. coli* consuming polysaccharide-derived sugars while locally depleting oxygen, thereby promoting oxygen-limited microenvironments.

While co-culture in microfluidics induces a broader spectrum of significantly regulated *B. thetaiotaomicron* genes as compared to co-culture in a flask, including several two-component systems (SI, Fig. S6, TCS in green), both liquid and microfluidic conditions promote induction of multiple polysaccharide utilization loci in the presence of *E. coli*, notably PUL77 (Fig. 1C) and PUL60 (Fig. 4C), consistent with diversified enzymatic expression under competitive conditions.

In the microfluidic device, co-culture was associated with changes in *E. coli* gene expression linked to aerobic respiration and central carbon oxidation (SI, Fig. S6B), indicating that spatially resolved oxygen availability is reflected at the transcriptional level. Consistent with a role for oxygen in shaping *E. coli* metabolism, *E. coli* grew markedly better in oxygenated *B. thetaiotaomicron* spent medium than under anoxic conditions, and this enhancement depended on starch-derived sugar uptake, as the Δ*pts* Δ*malE* mutant showed no growth increase (Fig. 3D). However, unlike well-mixed cultures where oxygen is continuously replenished, the microfluidic device imposes transport constraints under which oxygen consumption by *E. coli* can locally outpace its supply, increasing *B. thetaiotaomicron* benefit (SI, Fig. S4 and S7). This imbalance is expected to generate anoxic niches within the crypts, enabling a metabolic–environmental feedback that expands the habitat accessible to *B. thetaiotaomicron*.

### Metabolic and environmental feedback drive spatial organization and cooperative niche partitioning

To determine whether the spatial patterns observed in co-culture arise directly from *E. coli* ‘s ability to utilize starch-derived sugars and reshape the local oxygen environment, we compared the wild-type pairs with *B. thetaiotaomicron* co-cultured with the sugar-uptake–deficient *E. coli* Δ*pts* ΔmalE mutant. Under physiologically relevant oxygen gradients (0–0.5%), the species self-organized into complementary niches: *E. coli* localized along the oxygenated crypts (Fig. 5A, SI Fig. S8A). In contrast, abolishing *E. coli* sugar uptake reduced its growth along the crypt axis and limited *B. thetaiotaomicron* colonization of the distal region (Fig. 5B, SI Fig. S8B). In the absence of sugar-driven *E. coli* respiration, oxygen is expected to penetrate further into the crypt, reducing the spatial niche occupied by *B. thetaiotaomicron*. These results indicate that *E. coli*-mediated oxygen consumption contributes to maintaining the anoxic region that supports *B. thetaiotaomicron* colonization.

**Figure 5:**
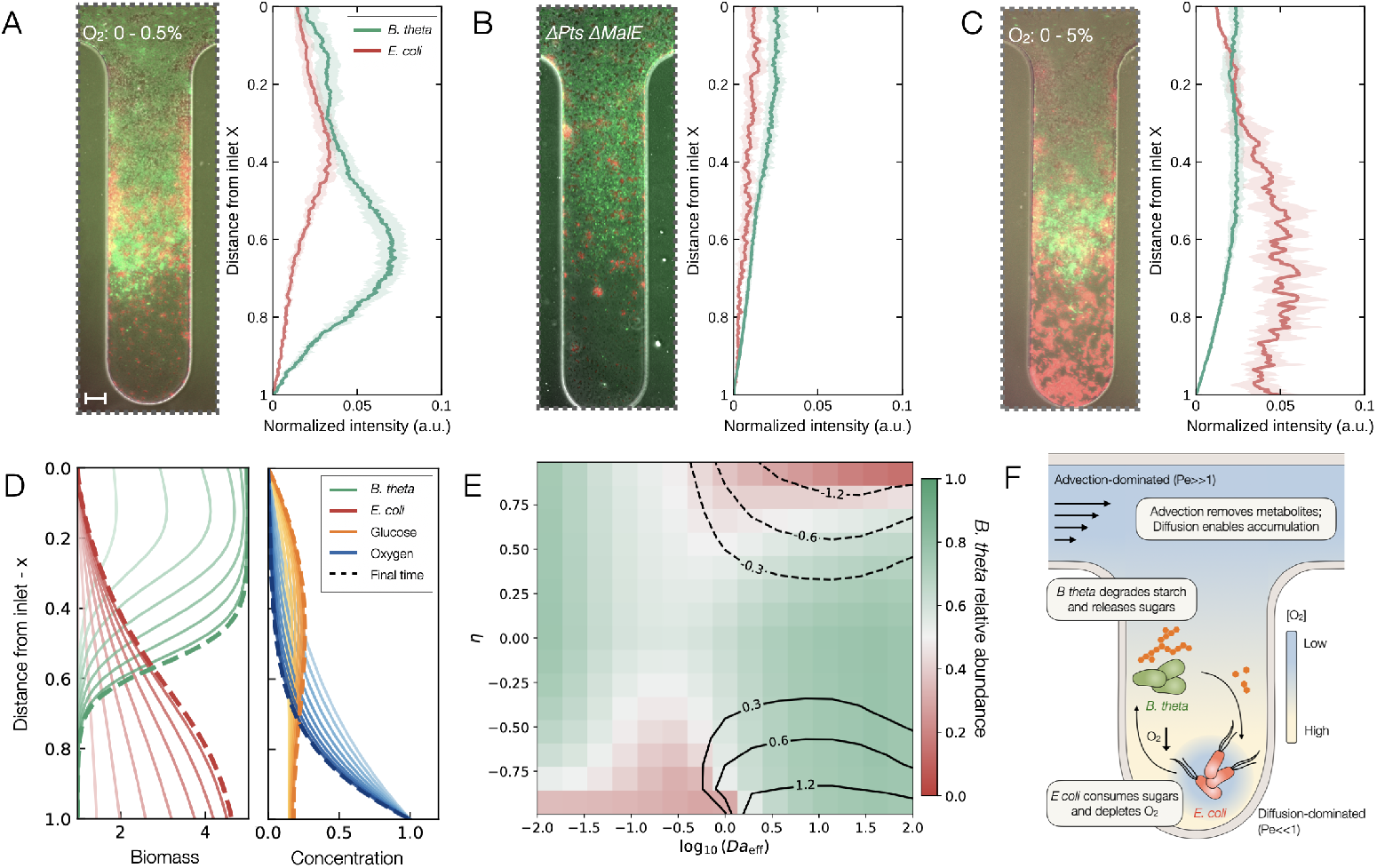
Spatial organization of *B. thetaiotaomicron* and *E. coli* along microfluidic crypts and corresponding reactive transport model. Fluorescence images and normalized intensity profiles showing the spatial distribution of *B. thetaiotaomicron* (green) and *E. coli* (red) along crypt depth under defined oxygen gradients. Profiles quantify species abundance as a function of distance from the inlet. Solid lines indicate averages (*n* = 3) and shaded areas standard deviation. Scale bar, 50 *µ*m. (A) Co-culture of wild-type strains under 0–0.5% oxygen. (B) Co-culture of *B. thetaiotaomicron* with the *E. coli* Δ*pts* Δ*malE* mutant under the same gradient. (C) Co-culture of wild-type strains under 0–5% oxygen. (D) Spatial profiles predicted by the one-dimensional reactive transport model along the crypt axis (0 = inlet, 1 = tip). Solid curves show temporal evolution of biomass and solute gradients; dashed lines indicate steady state. (E) Phase diagram of relative abundance as a function of environmental sensitivity (*η*) and effective Damköhler number (*Da*_eff_). Color indicates *B. thetaiotaomicron*/*E. coli* ratio; contour lines denote positive (solid) and negative (dashed) interactions, *I*_*B*_ = ln(*B*_coculture_*/B*_monoculture_). (F) Schematic of coupled transport and metabolic feedback. The main channel is advection-dominated (*Pe*≫1), while the cavity is diffusion-dominated (*Pe* ≪1), enabling solute accumulation. Oxygen diffuses from the cavity end and is depleted by *E. coli*, expanding the anoxic niche of *B. thetaiotaomicron*.

While the mucosa of the large intestine is typically hypoxic (*<* 1% O_2_) [30, 40, 41], oxygen availability can increase under conditions such as inflammation or post-antibiotic dysbiosis [28–30]. To test how elevated oxygen impacts spatial organization, we extended the gradient to 5%. Under these conditions, we observed a significant contraction of the *B. thetaiotaomicron* niche, while *E. coli* expanded throughout the entire crypt depth (Fig. 5C, SI Fig. S8C). While metabolic coupling is preserved across different oxygen gradients, the shift in relative abundance calls for a transition to a regime in which oxygen delivery outpaces microbial consumption. This suggests that spatial organization is not a static trait, but an emergent property governed by the competition between biological reaction kinetics and physical transport.

To capture this interplay mechanistically, we developed a one-dimensional reactive transport model that couples diffusion and consumption of oxygen, glucose, and amylopectin with logistic growth of each species under the experimental conditions. The model captures the essential metabolic–environment feedback: *B. thetaiotaomicron* colonizes anoxic regions where it degrades amylopectin and releases glucose; sugar accumulates in the crypt where it fuels *E. coli* growth; *E. coli*, in turn, consumes oxygen that diffuses from the crypt tip, widening the anoxic niche and enabling deeper *B. thetaiotaomicron* colonization than in monoculture (Fig. 5D, SI Fig. S9).

To understand how physical transport and metabolic feedback jointly shape species interactions, we explored the model across a range of conditions (Fig. 5E). Specifically, we varied two key model parameters: the ratio of reaction to transport time, quantified by the Damköhler number (*Da*), and the environmental sensitivity (*η*) of *B. thetaiotaomicron* fitness towards a generic compound (*O*). The Damköhler number reflects whether solutes are rapidly replenished by transport (low *Da*) or locally depleted by microbial activity (high *Da*). In parallel, *η* describes how the environment influences *B. thetaiotaomicron* growth: negative values characterize a hostile landscape where the compound *O* acts as a stressor (e.g., oxygen toxicity), while positive values characterize a landscape where the compound is an environmental requirement for reaching full carrying capacity. The absolute biomass benefit conferred by *E. coli* on *B. thetaiotaomicron* was largest at intermediate-to-high *Da* and inhibitory *η*: the regime where biological consumption outpaces physical delivery via metabolic feedback (Fig. 5E, SI Fig. S10). At low *Da*, transport dominates and the compound floods the cavity regardless of *E. coli* activity, leading to benefit collapse—consistent with the relative abundance recorded in well-mixed oxic liquid culture (SI Fig. S5). In contrast, when the compound effect is beneficial (*η >* 0), high *Da* promotes competitive interactions, as both species rely on the same compound, leading to a negative interaction (SI Fig. S11).

Together, these experimental and modeling results show that the interaction outcome is shaped by the interplay between physical transport and metabolic feedback (Fig. 5F). In well-mixed anoxic liquid cultures, both organisms are confined to a single shared niche, and *E. coli* behaves as a cheater—exploiting sugars released by *B. thetaiotaomicron* without reciprocal benefit. The microfluidic crypt fundamentally restructures this interaction by partitioning the environment into distinct microhabitats shaped by the joint action of transport and microbial metabolism. This spatial structure translates competitive exploitation into a self-organized division of labor: *B. thetaiotaomicron* degrades complex polysaccharides, supplying diffusible public goods, while *E. coli* locally depletes oxygen, expanding the anoxic niche on which *B. thetaiotaomicron* depends. Such emergent niche partitioning highlights how the balance between physical transport and metabolic activity influences microbial interaction outcomes.

## Discussion

A central challenge in microbial ecology is understanding how environmental spatial structure shapes microbial interactions [16, 42, 43]. Our results show that the physical environment is a key determinant of interaction outcomes. In well-mixed environments, polysaccharide-mediated interactions are dominated by competitive exploitation. The release of starch-derived sugars creates a globally accessible resource pool, allowing *E. coli* to benefit from the metabolic activity of *B. thetaiotaomicron* without providing reciprocal benefits. This is consistent with previous work showing that microbial interactions are typically dominated by competition and exploitation in environments where metabolites are broadly accessible [44, 45], supporting the view that widespread access to diffusible resources limits the emergence of cooperative interactions.

Introducing physical structure and transport reshapes the interaction motif by constraining spatial organization and redistributing secreted metabolites, thereby shifting the balance from competitive exploitation toward facilitation. In our system, crypt-like cavities promote the local accumulation of starch-derived sugars, generating spatially distinct growth opportunities. In contrast, in the flowing lumen, diffusible sugars are lost to advective washout, limiting the spread of exploitation across the system. This spatial partitioning limits uniform resource access by consumers and enables both species to occupy distinct ecological niches. Importantly, this organization depends on the metabolic substrate: replacing amylopectin with glucose abolishes spatial structure and results in a uniform competition. This is consistent with the localization of benefits from secreted compounds under constrained diffusion [46–50], as well as with the broader role of transport and spatial structure in shaping microbial interaction outcomes [25–27, 50–52].

Beyond nutrient localization, transport-induced gradients of oxygen throughout the crypt further shape the microbial interactions. *E. coli* respires oxygen, thereby expanding the anoxic niches of *B. thetaiotaomicron*. We show that this metabolic–environmental feedback emerges when transport and metabolic reaction timescales are balanced. This result emphasizes the importance of physical structure in mediating these feedbacks, which are increasingly recognized as key regulators of microbial interactions [12, 14, 53]. Our results suggest that while competitive exploitation dominates when metabolites are broadly accessible, the potential for cooperative interactions—arising from metabolic complementarity and contributing to community assembly [8, 54]—emerges under the physical partitioning of the environment. This mechanism is relevant in many natural environments, including host-associated systems, where oxygen gradients arise from physical constraints and biological activity. Consistent with previous works showing that changes in oxygen availability influence microbial composition [28–30], our results indicate that oxygen gradients modulate the spatial organization of metabolic interactions, and provide a mechanistic basis for how oxygen consumption by facultative anaerobes could contribute to the formation of anaerobic niches in the gut.

These environmental constraints are also reflected in the microbial transcriptome, revealing metabolic states associated with structured niches [55]. Co-culture in structured conditions elicited transcriptional responses consistent with substrate availability and local oxygen limitation. In *E. coli*, enhanced glucose metabolism and reduced aerobic respiration are consistent with growth under oxygen-limited conditions, whereas *B. thetaiotaomicron* modulates polysaccharide utilization loci in response to changes in metabolite availability. These responses are consistent with recent work showing that environmental gradients are reflected in localized gene expression patterns [56–58], supporting the view that microbial function depends on physical context.

Although our study focuses on a minimal two-species interaction, it captures a broader ecological motif in which polysaccharide degradation links primary degraders to different secondary consumers. Similar cross-feeding interactions involving *B. thetaiotaomicron* have been reported with opportunistic pathogens such as *Clostridioides difficile* [6] and *Salmonella Typhimurium* [59], as well as with other *Bacteroides* species [60, 61]. While this simplified system enables mechanistic resolution, natural microbial communities comprise many interacting taxa and additional environmental constraints that may modulate these interactions. Extending these approaches to more complex consortia will be important to determine how these mechanisms scale to community-level dynamics.

In conclusion, our findings show that microbial interaction outcomes emerge from the coupling between physical structure, metabolism and transport processes. Physical structure establishes the spatial context in which the chemical landscape is distributed, while microbial activity, in turn, reshapes local environmental conditions and creates new growth opportunities. Together, these findings provide a framework for understanding how the physical context and metabolic feedback shape microbial interaction outcomes across structured environments.

## Methods

### Bacterial strains and growth conditions

All cultures were grown at 37 °C in an anaerobic chamber (Coy Laboratory Products or Ruskinn anaerobic-microaerophilic station) under a gas mix of 2% H_2_ and 12% CO_2_ in N_2_. Frozen stocks of *Bacteroides thetaiotaomicron* VPI-5482 eGFP [62] and *E. coli* MG1655 Mars Red [63] were precultured overnight in BHI supplemented (BHIS) with Hemin 5 mg/L, NaCOH_3_ 2 gr/L, and cysteine 1 gr/L. Cells were collected by centrifugation (5 min, 5,000 ×*g*), washed in PBS, and inoculated at the desired OD into M9 defined medium. This consisted of M9 salts (6 gr/L Na_2_HPO_4_, 3 gr/L KH_2_PO_4_, 0.5 gr/L NaCl, 1 gr/L NH_4_Cl), 0.246 gr/L MgSO_4_.7H_2_O, 0.014 gr/L CaCl_2_.2H_2_O, 50 mg/Lt cysteine, 5 mg/L hemin, 2.5 *µ*g/L vitamin K3, 5 *µ*g/L vitamin B12, 2 mg/L FeSO_4_ 7H_2_O, and 1 gr/L carbon source (amylopectin, glucose, or maltose). *E. coli* S17*λ*pir was grown in Miller’s lysogeny broth (LB) (Corning) supplemented with ampicillin (100 *µ*g/ml) when required and incubated at 37 °C with 180 rpm shaking. Amylopectin from maize was autoclaved and dialysed using 3.5 kDa MW membranes (Slide-A-Lyzer Dialysis Cassettes, ThermoScientific). Amylopectin were autoclaved as 1% w/v in H_2_O and dialysed using 3.5 kDa MW membranes (Slide-A-Lyzer Dialysis Cassettes, ThermoScientific). Growth was assessed in liquid cultures inoculated at OD = 0.05, in 200 *µ*L volume wells by measuring OD_600_ using a plate reader (Tecan) with continuous orbital shaking. Colony-forming unit (CFU) was determined by serial dilution of cultures followed by spotting 5*µ*L drops of bacterial suspension on BHIS plates with selective antibiotics, erythromycin (15 *µ*g/mL) and kanamycin (25 *µ*g/mL) respectively for *B. thetaiotaomicron* and *E. coli*.

### Microfluidic fabrication

The microfluidic was fabricated using soft lithography [64]. We print the micromodel geometry into a silicon wafer via soft lithography, depositing a layer of SU-8 2150 (MicroChem Corp., Newton, MA) with controlled thickness (0.1 mm) via spin-coating. The wafer acts as a mold for liquid polydimethylsiloxane mixed with 10% by weight with its own curing agent (PDMS; Sylgard 184 Silicone Elastomer Kit, Dow Corning, Midland, MI). After solidification, we plasma-sealed the microfluidic device onto 25 mm × 75 mm glass slides.

### Microfluidic experiments and imaging

The geometry of the microfluidic device is characterized by a longitudinal main channel (width, W = 1 mm; height, H = 0.1 mm; length, L = 20 mm) flanked by crypt-like side cavities (width, w = 0.2 mm; height, h = 0.1 mm; length, l = 0.8 mm; interspace, i = 0.4 mm) (Fig. 4A). Medium was perfused through the central channel at a constant flow rate (Q), of 0.5 *µ*L/min with a syringe pump (PHD-ULTRA, Harvard Apparatus). The oxygen gradient along the orthogonal crypt-like cavities was controlled using a microscope-stage gas-mixing CO_2_–O_2_ incubator (PeCon). The module controlled the desired oxygen concentration on the microscope stage: 0.5% or 5%.

All equipments, buffers and media used in the microfluidic experiments were pre-equilibrated for 48 hours under an anoxic atmosphere in an anaerobic chamber (Coy Laboratory Products) under a gas mix of 2% H_2_ and 12% CO_2_ in N_2_. We used glass syringes (Hamilton) along with PTFE tubing and connectors, to ensure anoxic conditions in the central chip channel. We performed the microfluidic experiments within a constant temperature microscope incubator (OKOlab) maintained at 37 °C. Overnight bacterial cultures grown in BHIS were washed in PBS, pelleted, and then resuspended in M9 medium at OD = 0.01. First, we saturated the microfluidic device with PBS buffer and subsequently injected the bacterial suspensions in M9 medium, via the inlet port of the central chip channel. The entire chip volume was flushed twice with the bacterial suspension. The flow was then halted before initiating the perfusion of fresh, bacteria-free M9 medium for experiments lasting 24 h. After 24 hours, the microfluidic devices were exposed to atmospheric oxygen (~30 min were sufficient for full fluorescence recovery [62]) prior to quantify by fluorescence imaging eGFP and mars-Red proteins of the two strains. Imaging has been performed with inverted widefield microscopes (ScanR, Olympus or Eclipse Ti2, Nikon), equipped with a Hamamatsu ORCA flash 4.0 (16-bit, 6.5 *µ*m per pixel). We collected individual images (2048 × 2048) with a 10X magnification objective (0.65 *µ*m/pixel) in phase contrast and fluorescence optical configurations.

### ^13^C-labeled starch experiments

All cultures were grown at 37 °C in an anaerobic chamber (Coy Laboratory Products) under a gas mix of 2% H_2_ and 12% CO_2_ in N_2_. Frozen stocks of *Bacteroides thetaiotaomicron* VPI-5482 and *E. coli* MG1655 were precultured overnight in BHI supplemented with L-cysteine to 0.1% (w/v), and hemin to 0.5 mg/L. Cells were collected by centrifugation (5 min, 5,000 ×*g*), washed in PBS, and inoculated at OD_600_ 0.05 into M9 minimal medium supplemented with 50 mg/L L-cysteine, 5 mg/L hemin, 2.5 *µ*g/L vitamin K_1_, 2 mg/L FeSO_4_*·*7H_2_O, and 5 *µ*g/L vitamin B_12_. Cultures contained either 1 g/L of ^13^C starch or 1 g/L glucose. After 24 h of growth, cells were harvested (5 min, 5,000 ×*g*) and washed twice in PBS before cryo-CLsEM-FIB-SIMS analysis.

### Low-input bacterial RNA-seq

At the end of the experiments in liquid culture or in the microfluidic device, cells were collected in 200 *µ*L RNAprotect (Qiagen) to prevent RNA degradation, centrifuged, and the supernatant was removed. The resulting pellet was stored at −80 °C until extraction. Cell pellets were washed and resuspended in 100 *µ*l of PBS, and 1 *µ*l of a 1:20 dilution was dispensed in a 48-well plate pre-filled with 0.26 *µ*l PBS, 0.26 *µ*l lysis buffer (Takara), 0.1 U lysozyme (50 U/*µ*l; Epicentre), 0.03 *µ*l RNase inhibitor (100 U/*µ*l; Takara), and 1.95 *µ*l H_2_O. The plate was then sealed with an aluminum-based microfilm (Bio-Rad seal F), sonicated for 20 seconds (Sonorex Digitec DT 52, Bendelin), and spun down. Plates were kept at −80 °C until processing.

To generate cDNAs, a MATQ-Seq approach based on the protocol described in [35] was applied using the I.DOT dispensing robot (Dispendix). Briefly, reverse transcription was performed using Superscript IV (Invitrogen) and primers described in [65]. Next, all steps of the MATQ-Seq protocol (primer digestion, RNA digestion, poly(C) tailing, and second-strand synthesis) were performed as described in [35]. cDNA was purified using AMPure XP beads (Beckman Coulter) at a 1:1 (vol/vol) ratio. cDNA concentration was estimated using a Qubit Flex kit (Invitrogen), and quality was checked with a 2100 Bioanalyzer DNA High Sensitivity kit (Agilent Technologies).

All samples were treated in the same manner, independently of their composition. Library preparation was carried out using an I.DOT dispensing robot (Dispendix), and only one-tenth of the Nextera XT DNA library preparation kit (Illumina) was used compared to the manufacturer’s recommendations. Tagmentation was performed in a final volume of 2 *µ*l with 0.5 ng of cDNA input and a fragmentation time of 10 minutes. Next, index PCR was performed with 13 amplification cycles using the Nextera Index Kit v2 (set C). Libraries were purified with AMPure XP beads (Beckman Coulter) at a 1:1 (vol/vol) ratio, and their quality was assessed with a 2100 Bioanalyzer DNA High Sensitivity kit (Agilent Technologies).

Ribosomal RNA was depleted through a Cas9-based approach (DASH protocol [66]). Given the phylogenetic similarity between *E. coli* and *S. enterica*, oligo pools designed in [66] for *S. enterica* were used for rRNA depletion in *E. coli*. Generation of sgRNA pools and rRNA depletion were performed as described in [35], except for the following modifications. The Cas9:sgRNA complex was added to eight samples pooled equimolarly at a concentration of 2.1 fmol and a ratio of 1:2,000:2,000:2,000 (cDNA:Salmonella sgRNA pool:B. thetaiotaomicron sgRNA pool:Cas9). Digestion was performed by incubating the samples at 37 °C for 2 hours. Cas9 was deactivated by adding 1 *µ*l of Proteinase K (NEB) and incubating at 37 °C for 15 minutes. rRNA-depleted cDNA libraries were then purified through AMPure XP beads at a 1:1 (vol/vol) ratio and used as the input for a second PCR amplification (0.5 ng cDNA input) performed with the polymerase included in the Nextera XT kit (one-tenth of the recommended volume), using AWO-968 and AWO-969 as primers (complementary to the i5 and i7 indices, respectively) over 13 cycles. Cleanup was performed again with AMPure XP beads at a 1:1 (vol/vol) ratio. Pooled libraries were sequenced using an Illumina NextSeq P2 (100 cycles, single-end). After demultiplexing, reads were preprocessed, aligned, and quantified following the script described in [67]. For differential gene expression analysis, the edgeR package (version 4.2.1) was used with upper-quartile normalization. To correct for batch effects, sequencing data were further normalized using the RUVs correction method [68] with k = 3. Gene set enrichment analysis: The “fgseaRes” R package was used for pathway analysis with the following parameters minimum Size=10, maximum Size=500, eps=0, nproc= 1. Genes were assigned to functional categories based on their involvement in carbon source utilization, central carbon metabolism, and aerobic respiration. Functional annotations were derived from pathway information in EcoCyc [69] and complemented with regulatory information from RegulonDB [70]. Gene sets were manually curated based on these annotations. The complete list of genes and category assignments is provided in Supplementary Table S1. All raw and processed sequencing data have been deposited in the Gene Expression Omnibus (GEO) under accession number GSE327132.

### Construction of deletion mutants

Genetic constructions were made in *B. thetaiotaomicron* VPI-5482Δtdk background, developed for a two-step selection procedure of unmarked gene deletion by allelic exchange, as previously described [71]. All primers used in this study are listed below. Mutants were generated via allelic exchange using the suicide vector pLGB13 [72]. Construction of the mutant *B. thetaiotaomicron* ΔsusG was previously described in [73]. Approximately 500 bp regions flanking the target gene were amplified by PCR using Phusion Flash High-Fidelity PCR Master Mix (Thermo Fisher Scientific) and assembled with the plasmid backbone using Gibson assembly. The reaction mix, containing ISO buffer, T5 exonuclease, Phusion HF polymerase, and Taq DNA ligase, was incubated at 50 °C for 35 minutes. The assembled plasmid was then introduced into *E. coli* S17 *λ*pir, which served as the conjugation donor. For conjugation, exponentially growing cultures of donor and recipient were mixed at a 2:1 ratio and spotted on BHIS agar (Brain-Heart Infusion medium supplemented with 5 mg/L hemin, 2 g/L NaCOH3, and 1 g/L cysteine), followed by overnight incubation at 37 °C under aerobic conditions. The following day, the mix was plated on BHIS agar supplemented with erythromycin (15 *µ*g/mL) to select for *B. thetaiotaomicron* transconjugants that had undergone the first recombination, and gentamicin (200 *µ*g/mL) to inhibit donor *E. coli* growth. Resulting colonies were cultured overnight in antibiotic-free BHIS medium to facilitate plasmid loss, then plated on BHIS agar containing anhydrotetracycline to counterselect against cells retaining the vector. Candidate deletion mutants were verified by colony PCR using flanking primers, followed by Sanger sequencing.

The deletion of the operon ptsHI-crr in *E. coli* MG1655 was performed using lambda-red recombination [74]. The flippable version of a chloramphenicol resistance cassette was amplified by PCR (PCR master mix, Thermo Scientific, F548) using primers with long harms homologous to the flanking regions of ptsH and crr. The PCR product was then purified, dialyzed on a 0.025 *µ*m porosity filter and electroporated into a MG1655 strain bearing the pKOBEGA plasmid [75]. The resulting bacteria were spread on LB + Cm25 and incubated overnight at 37 °C. The obtained mutants were confirmed by PCR and Sanger sequencing and the absence of growth in presence of ampicillin confirmed that the cells expelled the pKOBEGA plasmid. The deletion was then transduced in the MG1655 WT background by P1vir phage transduction. The additional deletion of malE gene was then transduced from the keio collection strain JW3994 [76]. A list of all the primers used in this study can be found in SI Appendix, Table S2.

### Cryo-CLEM-FIB-SIMS

Vitrified samples were prepared by plunge-freezing as described previously [77]. Briefly, 3.5 *µ*l of sample (OD600 1-2) was applied to a freshly glow discharged 135 mesh Quantifoil London Finder grid (H15) and plunge-frozen into liquid ethane maintained at −178 °C, using a Vitrobot Mark IV at 10 °C and relative humidity of 95% after a wait time of 45 s, with −2 blot force, 0.5 s drain time and 2.5 s blot time.

The vitrified samples were subsequently imaged by cryo-CLEM-FIB-SIMS (correlated cryogenic light and electron microscopy with focused ion beam secondary ion mass spectrometry) for correlative spatial and chemical mapping [37]. The correlative imaging workflow encompasses three consecutive steps: cryo-light microscopy (cryo-LM) to distinguish the bacterial species via their fluorescent labelling, electron cryomicroscopy (cryo-EM) imaging yielding high spatial resolution images of the cells, and cryo-focused ion beam secondary ion mass spectrometry (cryo-FIB-SIMS) imaging providing spatially resolved chemical analysis [36], which allows the tracking of the 13C-labelled starch uptake of the respective bacterial species.

Fluorescence microscopy images were acquired under cryogenic conditions on a ZEISS LSM 900 confocal microscope equipped with an Airyscan 2 detector, Axiocam 306 camera, a Colibri 5 LED light source (ZEISS Microscopy) and a CMS196V3 (serial number DV5260-0009) cryo-stage (Linkam Scientific). Grid overview images were taken using a 0.5x/0.2 Numerical Aperture (NA) objective and high-resolution images were obtained using an LD EC Epiplan-Neofluar 100x/0.75 NA objective. Airyscan processing was performed for all high-resolution images in the Zen software (ZEISS Microscopy). Image processing was performed using Fiji [78].

Following the cryo-light microscopy imaging, the high spatial-resolution images of the vitrified bacterial cells were acquired using cryo-EM. Two-dimensional cryo-EM projection images were collected on a Titan Krios G1 microscope (ThermoFisher) operating at an acceleration voltage of 300 kV and equipped with a K3 direct electron detector (Gatan). Full cell images were recorded at nominal magnifications ranging from 6500x to 11500x, corresponding to pixel sizes of 23.89 to 18.01 Å, at exposure times of 2-5 s. Fourier bandpass filtering of the images was performed in Fiji [78].

Chemical mapping was performed by cryo-FIB-SIMS in a focused ion beam scanning electron microscope (FIB-SEM, Zeiss Crossbeam 550) equipped with a time-of-flight (ToF) mass spectrometer (TOFWERK fibTOF). During SIMS imaging, the sample is scanned by a focused gallium (Ga) ion beam. The secondary ions created during the interaction of ion beam and sample are extracted into the ToF mass spectrometer at every pixel, yielding a full mass spectrum per pixel [37]. Plotting the pixel-by-pixel intensity for each detected mass-to-charge ratio, allows the spatial visualisation of the chemical composition of the sample, where the colour scale represents the ion counts per extraction. The mass spectrometer can be operated in positive or negative ion mode, by respectively extracting either positive or negative secondary ions.

Cryo-FIB-SIMS imaging was performed at beam a voltage of 30 kV and beam current of 50 pA, the scanning area was 512 pixel by 512 pixel with mass spectrometry data collected using a binning of 8 pixel x 8 pixel, a FIB dwell time per pixel of 12.8 *µ*s, extraction pulse with of 1000 ns and magnification-dependent unbinned pixel sizes ranging from 10-30 nm. The mass resolution of the instrument is specified as *>* 700 by the manufacturer and was measured to be in the range of 1000-1500 in the experiments performed during this study.

### Reactive transport model

We developed a one-dimensional reactive transport model to describe the spatial dynamics of *Bacteroides thetaiotaomicron* (*B*) and *Escherichia coli* (*C*) and the associated gradients of glucose (*G*), oxygen (*O*), and amylopectin (*A*) along a cavity of length *L*.

Bacterial biomass densities evolve according to logistic growth:

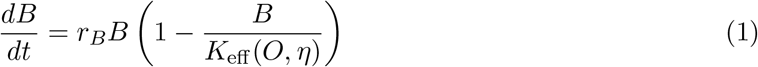

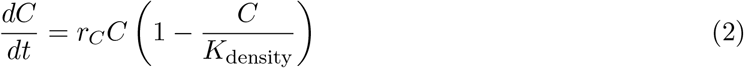

where *r*_*B*_ and *r*_*C*_ denote per-capita growth rates. The baseline carrying capacity is *K*_density_, while

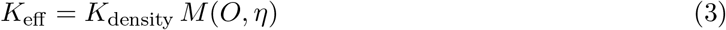

represents an oxygen-dependent effective carrying capacity for *Bacteroides thetaiotaomicron*, with *M* (*O, η*) a Hill-type response function. In the mechanistic simulations, *η* = −1, such that oxygen acts as an inhibitory factor for *Bacteroides thetaiotaomicron*.

Solute concentrations are governed by quasi-steady diffusion–reaction equations along the spatial coordinate *x*, expressed in nondimensional form. For example, the oxygen gradient is obtained from:

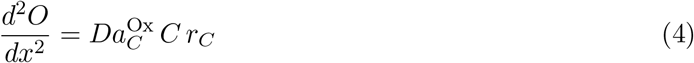

which couples diffusive transport to oxygen consumption by *Escherichia coli*. The Damköhler-like parameter *Da* scales the relative strength of biological reactions compared to diffusion, such that low values correspond to diffusion-dominated regimes, whereas high values lead to strong local depletion and steep spatial gradients. The same framework is applied to glucose and amylopectin.

At *x* = 0, concentrations are fixed such that *G* = *G*_0_, *O* = 0, and *A* = *A*_0_, while at *x* = *L*, zero-flux boundary conditions are imposed for glucose and amylopectin:

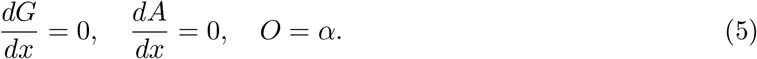

These conditions impose an anoxic-to-oxic gradient across the cavity, which is further reshaped by *Escherichia coli* respiration. Initial bacterial densities are spatially uniform, *B* = *C* = *ε*.

To explore the interplay between transport and biological activity, all reaction terms were scaled by an effective Damköhler factor *ϕ*, such that:

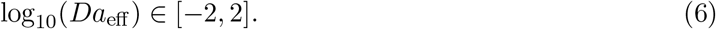

This formulation enables systematic exploration of regimes ranging from transport-dominated to reaction-dominated conditions.

To generalize this framework, we varied the environmental response parameter *η*, treating *O* as a generic compound whose effect on *Bacteroides thetaiotaomicron* can be inhibitory (*η <* 0), neutral (*η* = 0), or beneficial (*η >* 0). This enables mapping of interaction outcomes across the (*Da*_eff_, *η*) parameter space.

Spatial solute fields were solved quasi-steadily at each biomass update step using a boundary value solver. Model outputs included total biomass, relative abundance, oxygen depletion, and interaction strength, defined as:

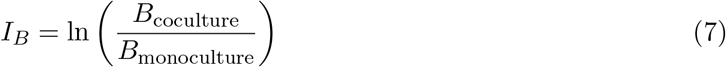

where biomass was integrated over time.

A full mathematical description, including governing equations, parameter definitions, and numerical implementation, is provided in the Supplementary Information and in Table S3.

### Transport simulation

Fluid flow velocity and oxygen transport through the microfluidic chip were simulated in COMSOL Multiphysics by coupling the *Laminar Flow* and *Transport of Diluted Species* modules. The geometry comprised a PDMS membrane (thickness ~ 0.01 mm) that separated the lateral channels filled with controlled gas from the central fluid channel (height ~ 0.10 mm) filled with culture medium.

Fluid flow was described by the steady-state incompressible Navier–Stokes equations, while oxygen transport was modeled using Fick’s laws of diffusion, accounting for both diffusion and advection in the channel and diffusion across the PDMS layer.

Oxygen diffusivities and solubilities for PDMS (*D*_PDMS_ ≈ 3.25 × 10^−5^ cm^2^/s, *S*_PDMS_ ≈ 1.62 mmol L^−1^ atm^−1^) and medium (*D*_medium_ ≈ 1.9 × 10^−5^ cm^2^/s, *S*_medium_ ≈ 0.22 mmol L^−1^ atm^−1^) were set to previously measured values [79].

Boundary conditions included a fixed oxygen concentration at the PDMS outer surface and zero-flux at symmetry planes or device walls. Mesh convergence was verified, and steady-state and transient simulations were used to predict oxygen gradients across the fluid channel and PDMS layer. The resulting oxygen concentration profiles were used to estimate the gradient magnitude (e.g., 0–0.5% O_2_) and its spatial extent.

### Statistics

Statistical analysis was performed in RStudio (v 2024.12.0+467). Unless otherwise indicated, results represent the mean *±* standard deviation. Statistical significance is indicated as \**P <* 0.05, \*\**P <* 0.01, and \*\*\**P <* 0.001.

## Acknowledgements

We thank Sylvie Létoffé, Sol Vendrell-Fernández and Yutaka Yoshii for their contribution to mutant design, and Minhee Kim and Julia Bos for their help.

This work was funded by the Swiss National Science Foundation Grants (P500PB_211100 and P5R5-3_235220, D.S.); Institut Pasteur and grants from the French government’s Investissement d’Avenir Program, Laboratoire d’Excellence “Integrative Biology of Emerging Infectious Diseases” (grant n°ANR-10-LABX-62-IBEID to J.M.G), the Medical Research Council, UK (MC UP 1201/31 to TAMB), the Wellcome Trust (225317/Z/22/Z to TAMB), and the Leverhulme Trust (Philip Leverhulme Prize to TAMB). K.R.P. acknowledges funding by ERC (project-ID # 866028). A.J.W. acknowledges funding by the European Research Council (ERC) (project-ID #101040214) and by the German Research Foundation (DFG) through the SFB1583 DECIDE (#492620490; project A04) and the Cluster for Nucleic Acid Sciences and Technologies – NUCLEATE (project-ID 533767322 – EXC 3113/1).

## Author contributions

D.S. and J.M.G. conceived the study. D.S. designed and performed the microfluidic experiments. E.B. conducted RNA-sequencing. E.B., D.S., and L.D. analyzed the transcriptomic data. S.B. contributed to the microbiological assays. K.R.P. contributed to the design of isotope labeling experiments. H.O. and T.B. carried out Cryo-CLEM-FIB-SIMS analyses. D.S., S.T.R., and C.B. designed the knockout mutants. Y.D. and D.S. performed numerical simulations. D.S. wrote the original manuscript draft with J.M.G. and A.J.W.; all authors reviewed the final manuscript.

