## Supplementary Information for "Oxygen gradients reshape cross-feeding through emergent spatial organization of gut commensal bacteria"

[1]Institut Pasteur, Université Paris Cité, CNRS UMR 6047, Genetics of Biofilms laboratory, Paris F-75015, France
[2]Medical Research Council Toxicology Unit, University of Cambridge, UK
[3]Department of Microbiology, Biocenter, University of Würzburg, D-97074, Würzburg, Germany
[4]Helmholtz Institute for RNA-based Infection Research (HIRI), Helmholtz Centre for Infection Research (HZI), Würzburg, D-97080, Germany
[5]Cluster for Nucleic Acid Sciences and Technologies – NUCLEATE, Würzburg, D-97080, Germany
[6]Structural Studies Division, MRC Laboratory for Molecular Biology, Cambridge, UK
[7]Institut de Mécanique des Fluides de Toulouse, Université de Toulouse, CNRS UMR 5502, France
[8]Institut Pasteur, Université Paris Cité, Biomaterials and Microfluidics core facility, Paris F-75015, France

### I. Supplementary Figures

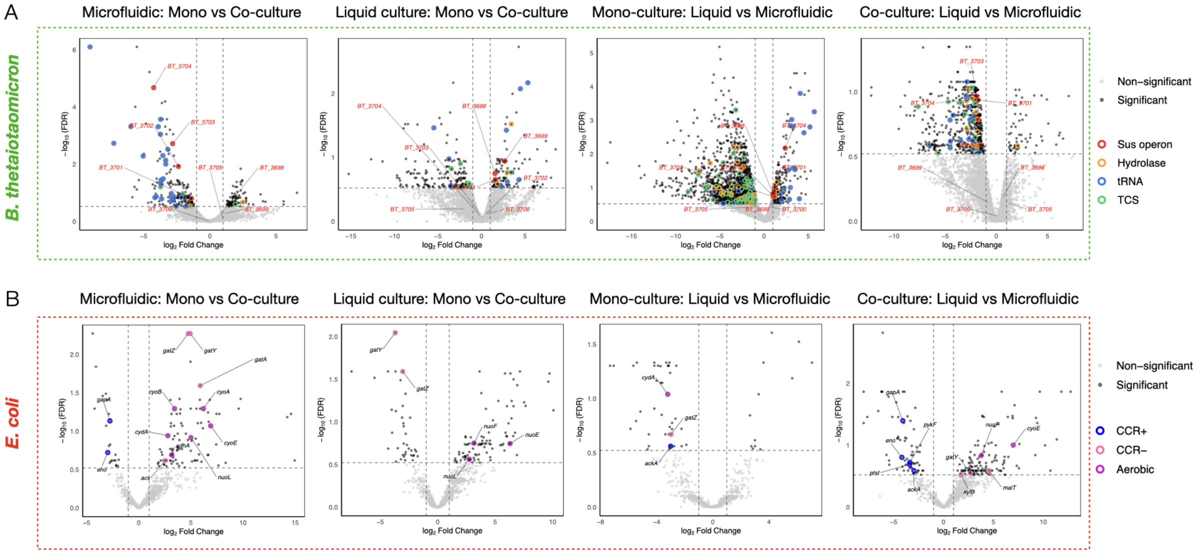


**Figure S1: Transcriptional responses to culture mode and growth environment in *Bacteroides thetaiotaomicron* and *Escherichia coli*.** Volcano plots showing differential gene expression across mono- and co-culture conditions in microfluidic devices and liquid culture. (A) For *Bacteroides thetaiotaomicron*, comparisons include mono- versus co-culture in microfluidic devices, mono- versus co-culture in liquid culture, and liquid culture versus microfluidic growth in both mono- and co-culture conditions, as indicated above each panel. (B) The same set of comparisons is shown for *Escherichia coli*. The x-axis shows $\log_2$ fold change and the y-axis shows $-\log_{10}$ false discovery rate (FDR). Genes with $\log_2$ fold change $\geq 1$ and FDR $\leq 0.3$ are highlighted. In *Bacteroides thetaiotaomicron*, genes from the starch polysaccharide utilization locus (Sus operon; BT_3698–BT_3705) are highlighted in red, Sus-associated homologs in green, tRNA genes in blue, hydrolases in orange, and two-component systems in cyan. In *Escherichia coli*, highlighted genes correspond to CCR-positive, CCR-negative, and aerobic metabolism-associated categories.

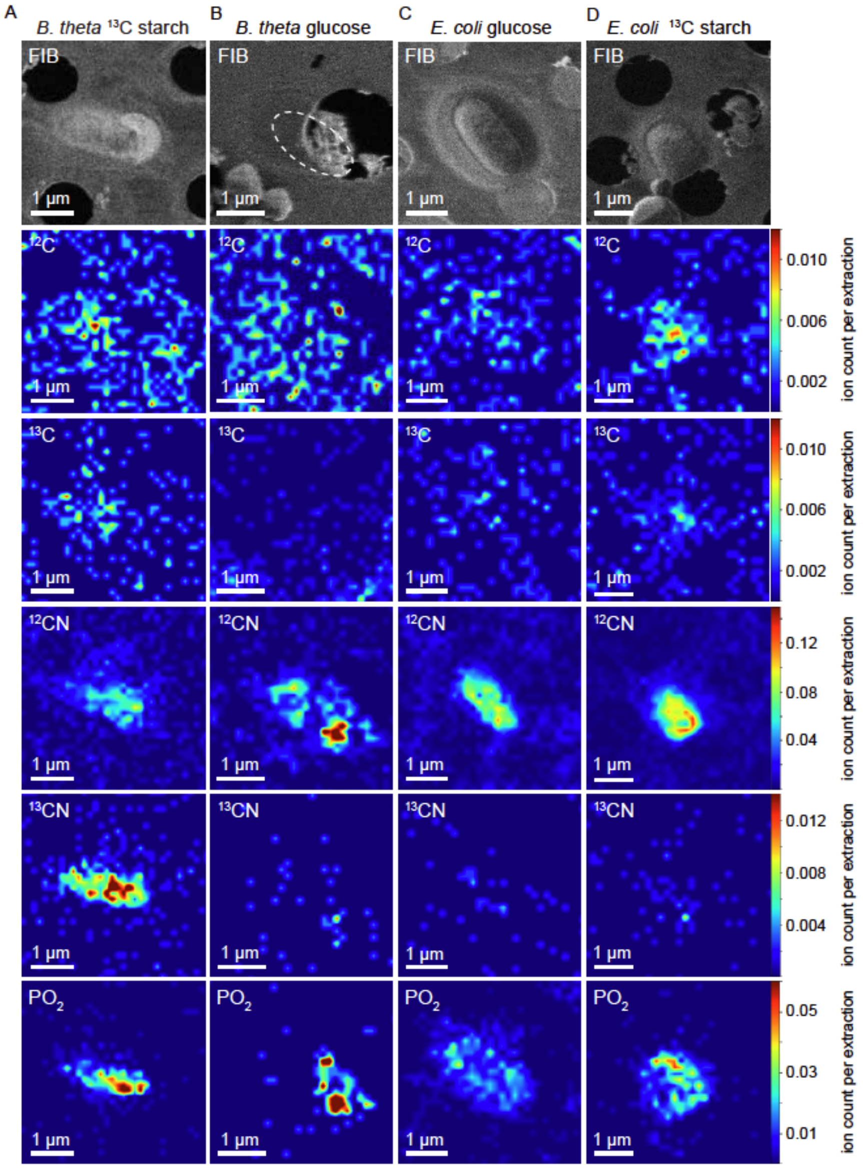


**Figure S2: SIMS data for single-culture control experiments.** Secondary electron FIB image and cryo-FIB-SIMS images for $^{12}C$, $^{13}C$, $^{12}CN$, $^{13}CN$, and $PO_2$ for cultures of *Bacteroides thetaiotaomicron* in medium supplemented with $^{13}C$-labelled starch. (A) *Bacteroides thetaiotaomicron* grown in unlabelled medium with glucose as the carbon source. (B) *Escherichia coli* grown in unlabelled medium with glucose as the carbon source. (C) *Escherichia coli* grown in medium supplemented with $^{13}C$-labelled starch. Secondary electron FIB images are provided to clarify the spatial localization of the cells. SIMS images for each ionic species were scaled to the same colour scale to facilitate comparison.

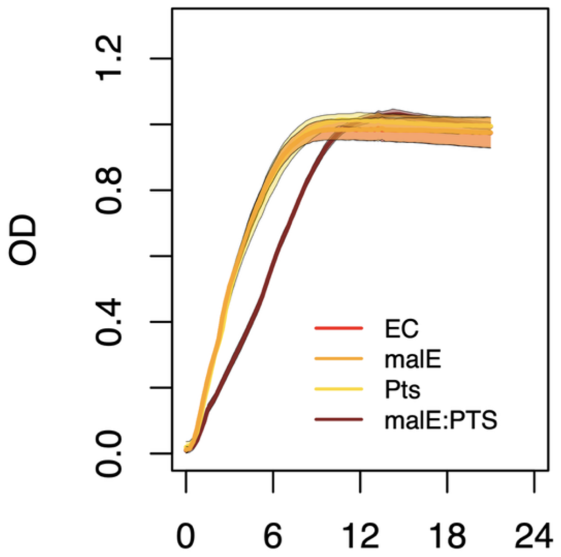


**Figure S3: Sugar uptake mutants of *Escherichia coli* display normal growth in rich medium.** Growth curves of *Escherichia coli* wild type (WT), $\Delta pts$ (glucose-PTS deficient), $\Delta malE$ (maltose transport deficient), and $\Delta pts\Delta malE$ mutants cultured in oxic LB medium. Optical density ($OD_{600}$) was monitored over time.

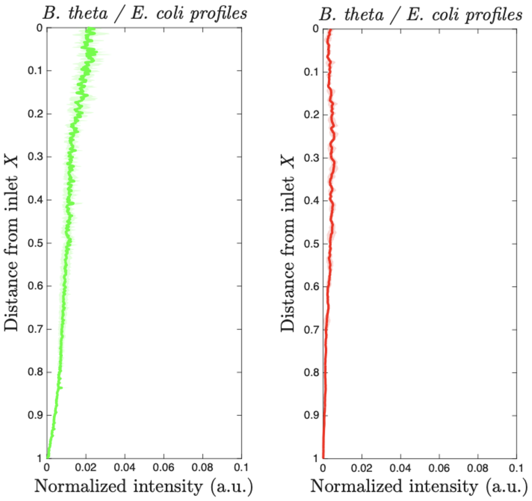


**Figure S4: Biomass profiles of *Bacteroides thetaiotaomicron* and *Escherichia coli* monocultures in microfluidic crypts under oxygen gradients.** Spatial biomass distributions of *Bacteroides thetaiotaomicron* (left) and *Escherichia coli* (right) monocultures grown for 24 h in the microfluidic device under an imposed oxygen gradient spanning 0–0.5% $O_2$ along the crypt axis. Fluorescence intensity was quantified along the cavity length and normalized for each strain. *Bacteroides thetaiotaomicron* preferentially accumulated in oxygen-limited regions, whereas *Escherichia coli* exhibited only minimal background growth.

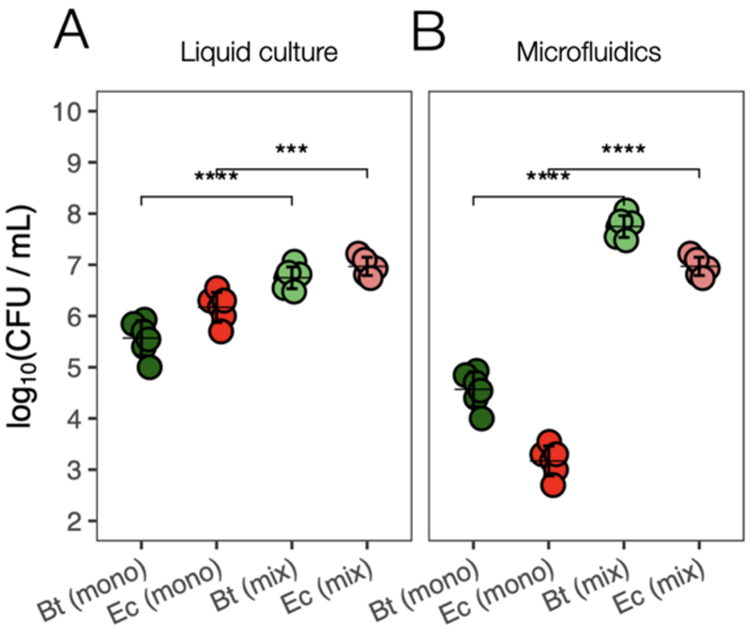


**Figure S5: Spatial structure alters competitive outcomes between *Bacteroides thetaiotaomicron* and *Escherichia coli*.** Colony-forming unit (CFU) counts measured after 24 h growth of monocultures and cocultures of *Bacteroides thetaiotaomicron* and *Escherichia coli* in (A) liquid culture exposed to oxygen and (B) the microfluidic device under a stable oxygen gradient (0–0.5%).

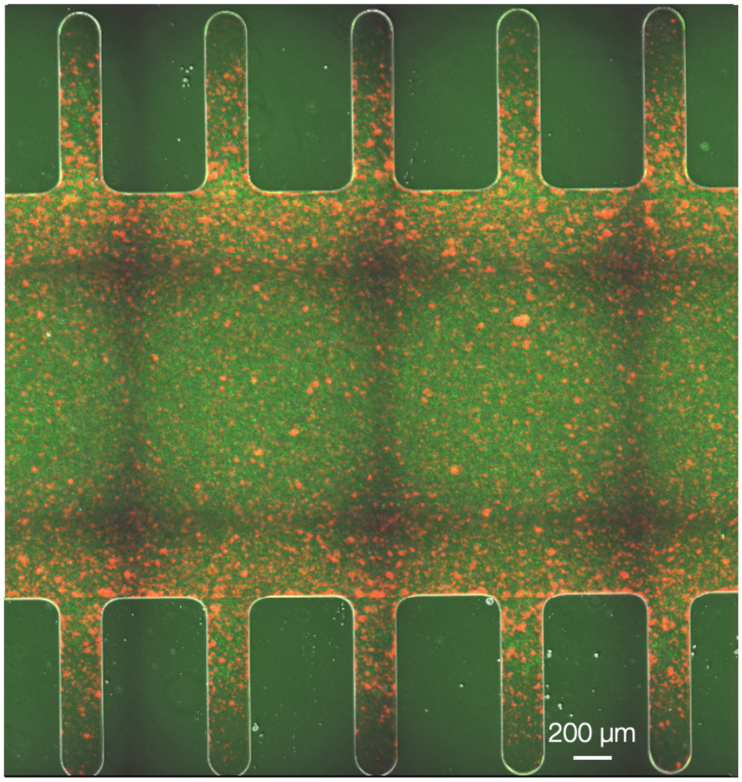


**Figure S6: Uniform glucose availability abolishes spatial segregation in the microfluidic device.** Fluorescence micrographs showing the spatial distribution of *Escherichia coli* (red) and *Bacteroides thetaiotaomicron* (green) grown in the microfluidic device with M9 medium supplemented with glucose as the sole carbon source after 24 h. Under these conditions, both species display more homogeneous spatial distributions along the crypt axis, indicating that spatial organization observed with amylopectin depends on polysaccharide-mediated cross-feeding.

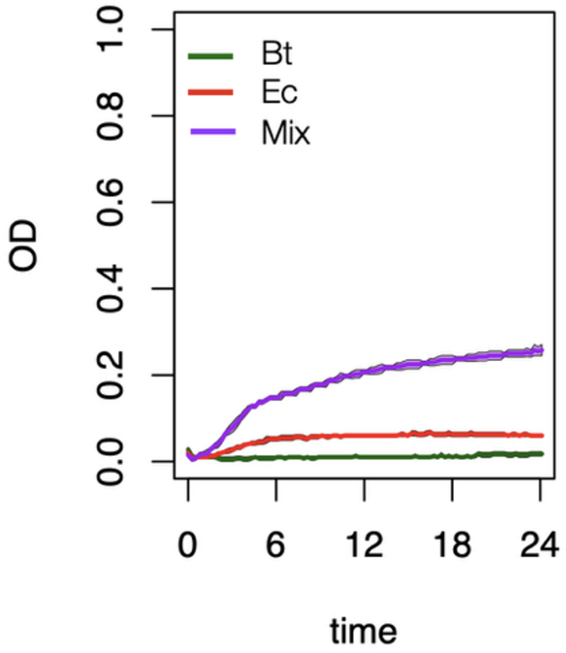


**Figure S7: Oxygen presence alters coculture dynamics in liquid medium.** Growth curves of *Bacteroides thetaiotaomicron* and *Escherichia coli* grown as monocultures or cocultures in M9 medium supplemented with amylopectin under oxic conditions. Optical density ($OD_{600}$) was monitored over 24 h.

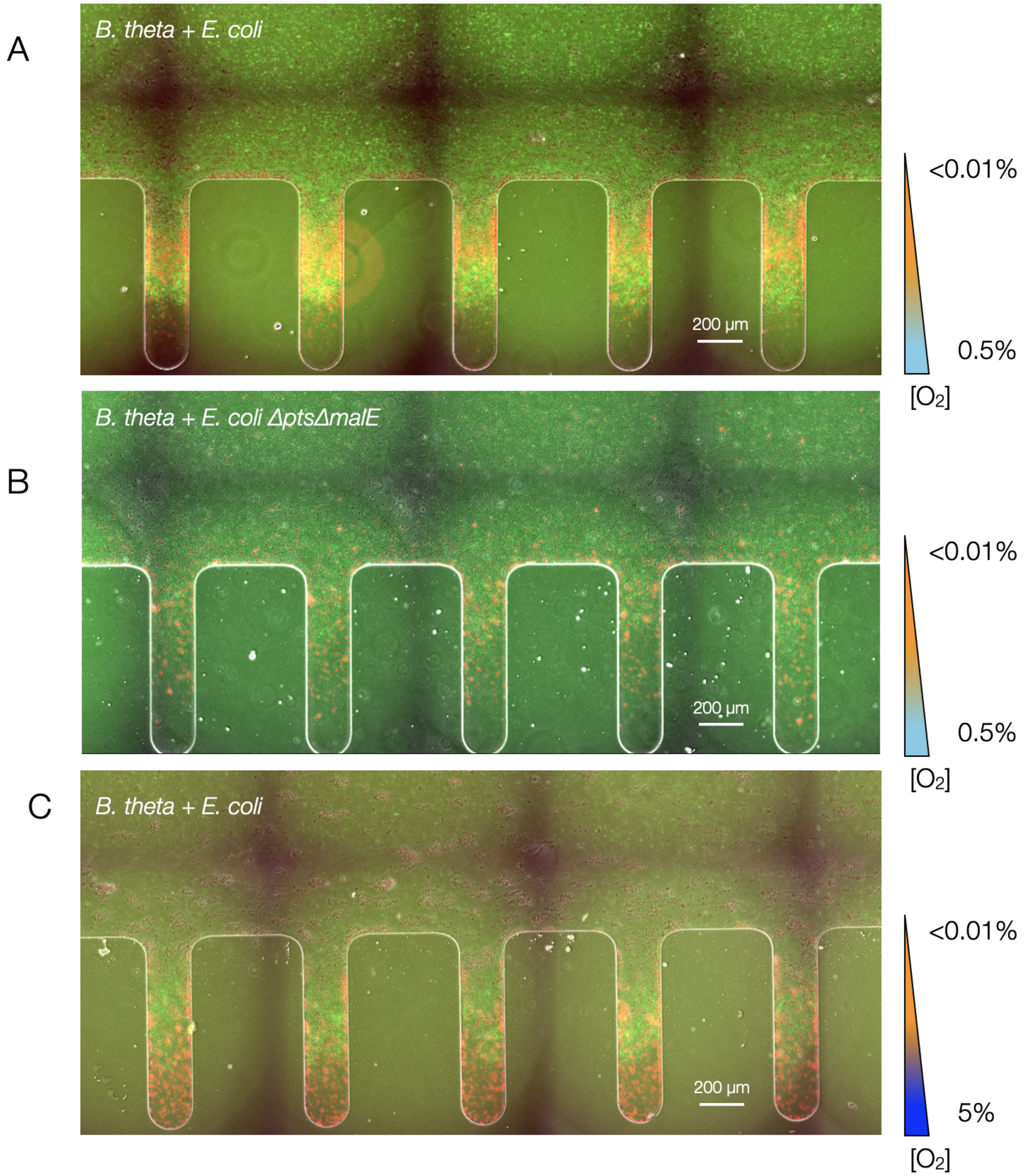


**Figure S8: Large-scale spatial organization of cocultures in different conditions.** Wide-field fluorescence images showing spatial organization of *Bacteroides thetaiotaomicron* (green) and *Escherichia coli* (red) after 24 h growth in the microfluidic device. (A) Coculture of *Bacteroides thetaiotaomicron* with *Escherichia coli* WT under a 0–0.5% $O_2$ gradient. (B) Coculture of *Bacteroides thetaiotaomicron* with *Escherichia coli* $\Delta pts \Delta malE$ under the same oxygen gradient. (C) Coculture of *Bacteroides thetaiotaomicron* with *Escherichia coli* WT under an expanded oxygen gradient (0–5%). Color overlays indicate local oxygen concentrations. Scale bars, 200 $\mu$m. Disruption of sugar uptake in *Escherichia coli* or increased oxygen availability alters the emergent spatial organization.

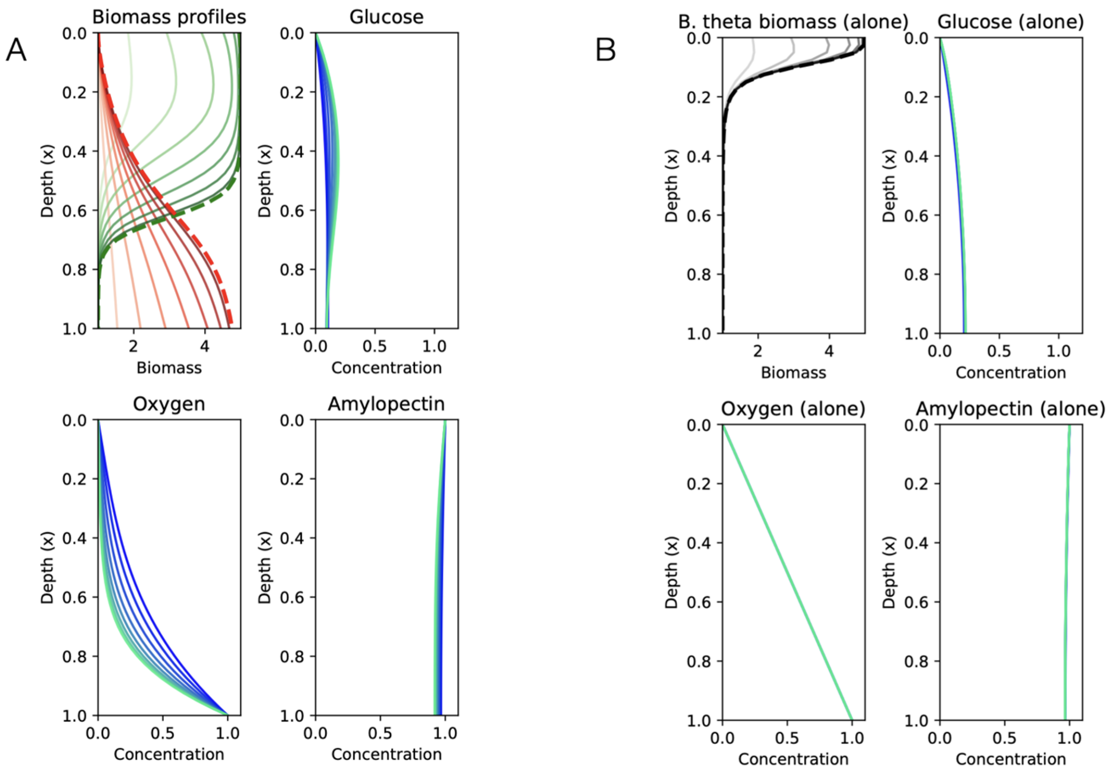


**Figure S9: Temporal evolution of spatial gradients in co-culture and monoculture.** Spatial profiles of biomass and metabolites along the depth axis are shown over time for (A) co-culture and (B) *Bacteroides thetaiotaomicron* monoculture at $\eta = -0.8$ and $\log_{10}(Da_{\text{eff}}) = 0$. In co-culture, *Escherichia coli* growth and oxygen consumption generate steep oxygen gradients, leading to spatially structured biomass distributions and localized depletion of glucose. In monoculture, profiles exhibit weaker gradients and more uniform metabolite distributions.

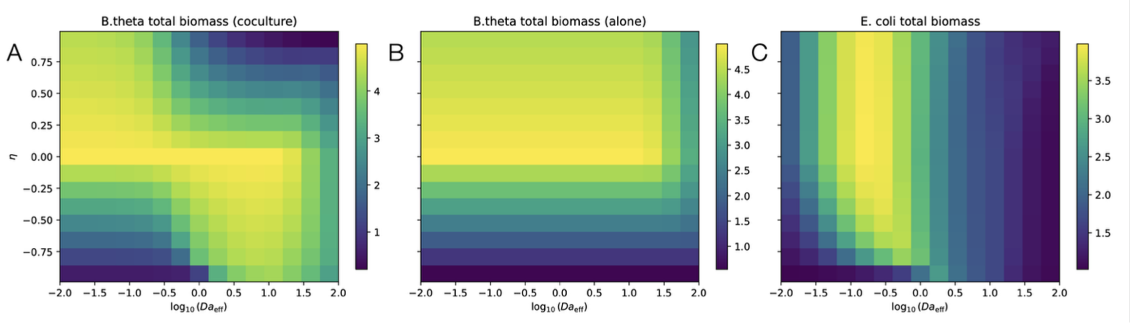


**Figure S10: Total biomass across transport regimes and interaction parameters.** Total biomass of (A) *Bacteroides thetaiotaomicron* in co-culture, (B) *Bacteroides thetaiotaomicron* in monoculture, and (C) *Escherichia coli* in co-culture, shown as a function of environmental sensitivity ($\eta$) and the effective Damköhler number ($Da_{\text{eff}}$).

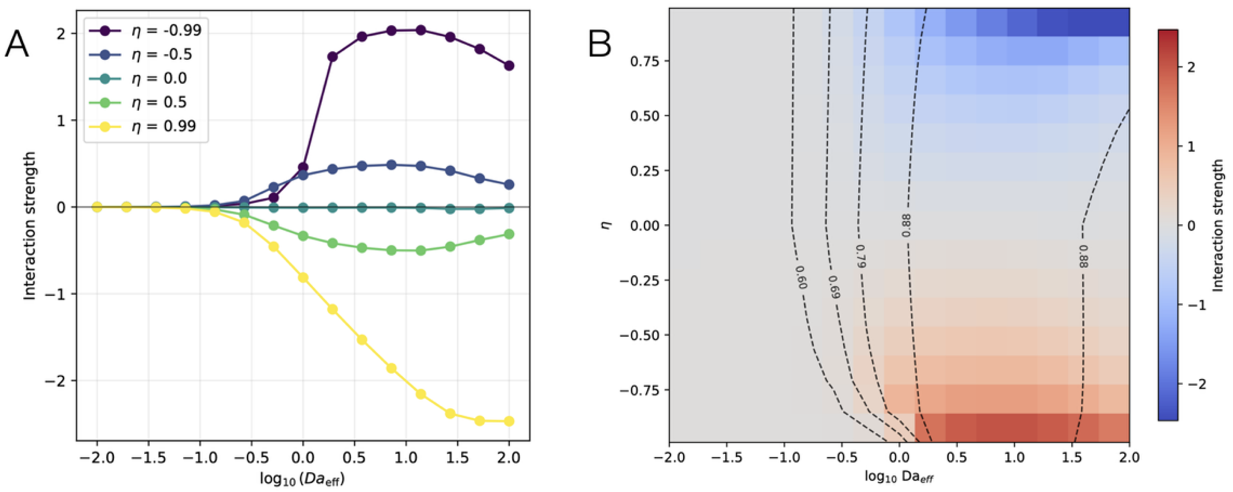


**Figure S11: Interaction strength across transport regimes and environmental sensitivity.** Interaction strength $I = \ln(B_{\text{theta,co}}/B_{\text{theta,mono}})$. (A) Interaction strength as a function of $Da_{\text{eff}}$ for selected $\eta$ values. (B) Interaction strength across the full parameter space of $\eta$ and $\log_{10}(Da_{\text{eff}})$. Positive and negative values indicate facilitative and inhibitory interactions, respectively. Contour lines indicate the fraction of compound (oxygen) removed by *Escherichia coli*.

#### II. Reactive–transport model

The model represents a dead-end cavity connected to a flowing channel, analogous to a colonic crypt or a stagnant mucus layer. The main channel is advection-dominated, whereas the cavity is diffusion-dominated due to negligible flow. We therefore restrict the model to the cavity and neglect advection.

The relevant transport regime is characterized by the Péclet number:

$$\text{Pe} = \frac{UL}{D}, \tag{1}$$

where $U$ is the flow velocity and $D$ the solute diffusivity. In the cavity, $\text{Pe} \ll 1$ (typically near zero in mucus-filled geometries), justifying a purely diffusive description. The spatial coordinate is normalized such that $x \in [0, 1]$.

The system consists of:

$$B(x,t), \quad C(x,t) \quad \text{(biomass densities)} \tag{2}$$

$$G(x), \quad O(x), \quad A(x) \quad \text{(solute concentrations)} \tag{3}$$

Solute concentrations are non-dimensionalized by their boundary values $(G_0, A_0, \alpha)$, while biomass remains dimensional to maintain consistency with the density-dependent carrying capacity. Solute fields are treated as quasi-steady (see below), while biomass evolves dynamically.

To model a wide range of biological responses to the environment, we define a sensing function $S(O)$, that represents the microbial perception of the compound. We utilize a Hill coefficient $(n)$

to characterize the steepness of the sensing threshold:

$$S(O) = \frac{O^n}{O^n + K_{\text{env}}^n}. \tag{4}$$

The impact of this compound on the species is governed by the environmental response parameter $\eta$, which determines the modifier function $M(O, \eta)$:

$$M(O, \eta) = \begin{cases} 1 + \eta S(O), & \eta \leq 0, \\ 1 - \eta(1 - S(O)), & \eta > 0. \end{cases} \tag{5}$$

Biomass evolves according to logistic growth:

$$\frac{\partial B}{\partial t} = r_B B \left(1 - \frac{B}{K_{\text{eff}}(O, \eta)}\right), \tag{6}$$

$$\frac{\partial C}{\partial t} = r_C C \left(1 - \frac{C}{K_{\text{density}}}\right). \tag{7}$$

The carrying capacity of species $C$ is constant, while that of species $B$ depends on the compound ($O$) through an effective carrying capacity:

$$K_{\text{eff}} = K_{\text{density}} M(O, \eta). \tag{8}$$

Growth follows Monod kinetics. For species $B$:

$$r_B = r_{B,\max}^{\text{Gluc}} \frac{G}{G + K_B^{\text{Gluc}}} + r_{B,\max}^{\text{Am}} \frac{A}{A + K_B^{\text{Am}}}. \tag{9}$$

For species $C$, growth is modeled to include both aerobic and anaerobic components, reflecting its facultative nature:

$$r_C = r_{C,\max} \left(\frac{G}{G + K_C^{\text{Gluc}}}\right) \left[\frac{O}{O + K_C^{\text{Ox}}} + f_{\text{ana}} \left(1 - \frac{O}{O + K_C^{\text{Ox}}}\right)\right]. \tag{10}$$

Solutes satisfy quasi-steady diffusion–reaction equations:

Glucose:

$$\frac{d^2 G}{dx^2} = Da_B^{\text{Gluc}} B r_B^{\text{Gluc}} + Da_C^{\text{Gluc}} C r_C - \lambda Pa_B^{\text{Gluc}} B r_B^{\text{Am}}, \tag{11}$$

Environmental compound ($O$):

$$\frac{d^2 O}{dx^2} = Da_C^{\text{Ox}} C r_C^{\text{aer}}, \tag{12}$$

Amylopectin:

$$\frac{d^2 A}{dx^2} = Da_B^{\text{Am}} B r_B^{\text{Am}}. \tag{13}$$

We assume:

$$\frac{\partial G}{\partial t} \approx 0, \quad \frac{\partial O}{\partial t} \approx 0, \quad \frac{\partial A}{\partial t} \approx 0. \tag{14}$$

Reaction–transport coupling is expressed through effective Damköhler numbers:

$$Da = \frac{\text{reaction rate scale}}{\text{diffusion scale}}. \tag{15}$$

All reaction terms are scaled by a common factor $\phi$:

$$Da_i = \phi Da_{i,0}. \tag{16}$$

We define $Da_{\text{eff}} \propto \phi$.
At $x = 0$:

$$G = G_0, \quad O = 0, \quad A = A_0, \tag{17}$$

At $x = L$:

$$\frac{dG}{dx} = 0, \quad O = \alpha, \quad \frac{dA}{dx} = 0. \tag{18}$$

At each time step:

1. Solve steady diffusion–reaction equations for $(G, O, A)$.
2. Compute local growth rates.
3. Update biomass using explicit time stepping.

The boundary value problem is solved using a collocation method (`bvp4c` in MATLAB).
Total biomass:

$$B_{\text{tot}} = \int_0^L B(x)\, dx. \tag{19}$$

Relative abundance:

$$f_B = \frac{B_{\text{tot}}}{B_{\text{tot}} + C_{\text{tot}}}. \tag{20}$$

Interaction strength:

$$I_B = \ln\left(\frac{\int B_{\text{tot}}^{\text{co}}(t)\, dt}{\int B_{\text{tot}}^{\text{mono}}(t)\, dt}\right). \tag{21}$$

All model parameters are listed in Table S3.

**Table S1: List of *Escherichia coli* genes involved in carbon source utilization and aerobic metabolism.**

| Gene | Category | Gene | Category | Gene | Category |
|---|---|---|---|---|---|
| *lamB* | CCR− | *crr* | CCR+ | *cyoA* | Aerobic |
| *malE* | CCR− | *ptsH* | CCR+ | *cyoB* | Aerobic |
| *malF* | CCR− | *ptsI* | CCR+ | *cyoC* | Aerobic |
| *malG* | CCR− | *manX* | CCR+ | *cyoD* | Aerobic |
| *malK* | CCR− | *manY* | CCR+ | *cyoE* | Aerobic |
| *malS* | CCR− | *manZ* | CCR+ | *cydA* | Aerobic |
| *malP* | CCR− | *galP* | CCR+ | *cydB* | Aerobic |
| *malQ* | CCR− | *glk* | CCR+ | *nuoA* | Aerobic |
| *malZ* | CCR− | *pgi* | CCR+ | *nuoB* | Aerobic |
| *malM* | CCR− | *pfkA* | CCR+ | *nuoC* | Aerobic |
| *malT* | CCR− | *fbaA* | CCR+ | *nuoD* | Aerobic |
| *malX* | CCR− | *gapA* | CCR+ | *nuoE* | Aerobic |
| *malY* | CCR− | *pgk* | CCR+ | *nuoF* | Aerobic |
| *gatA* | CCR− | *gpmA* | CCR+ | *nuoG* | Aerobic |
| *gatB* | CCR− | *eno* | CCR+ | *nuoH* | Aerobic |
| *gatC* | CCR− | *pykF* | CCR+ | *nuoI* | Aerobic |
| *gatD* | CCR− | *pykA* | CCR+ | *nuoJ* | Aerobic |
| *gatY* | CCR− | *ackA* | CCR+ | *nuoK* | Aerobic |
| *gatZ* | CCR− | *pta* | CCR+ | *nuoL* | Aerobic |
| *mglA* | CCR− | | | *nuoM* | Aerobic |
| *mglB* | CCR− | | | *nuoN* | Aerobic |
| *mglC* | CCR− | | | *sdhA* | Aerobic |
| *xylA* | CCR− | | | *sdhB* | Aerobic |
| *xylB* | CCR− | | | *sdhC* | Aerobic |
| *rhaA* | CCR− | | | *sdhD* | Aerobic |
| *rhaB* | CCR− | | | | |
| *rhaD* | CCR− | | | | |
| *acs* | CCR− | | | | |
| *araA* | CCR− | | | | |
| *araB* | CCR− | | | | |
| *araD* | CCR− | | | | |
| *mtlA* | CCR− | | | | |
| *mtlD* | CCR− | | | | |
| *srlA* | CCR− | | | | |
| *srlB* | CCR− | | | | |
| *srlE* | CCR− | | | | |

#### Table S2: List of primers used in this study.

| Construct | Name | Sequence (5'→3') |
|---|---|---|
| pLGB13 | pLGB13-linF | gcttatcgataccgtcgac |
| | pLGB13-linR | tgatatcgaattcctgcagc |
| | pLGB13-chF | GGTGTAAGATTAGCATTATGAGTG |
| | pLGB13-chR | CCATCACTGGAAGATAGGC |
| Δ*susG* (BT_3698) | 3698-5F | GCTGCAGGAATTCGATATCAgggataaagatatgatgagtgaggaag |
| | 3698-5R | catcatcattatgaataaacatctccacttcaagttgggcaactaaatac |
| | 3698-3F | gtatttagttgcccaacttgaagtggagatgtttattcataatgatgatg |
| | 3698-3R | GTCGACGGTATCGATAAGCgctctcgatgaaaacctgtatatactggaagc |
| Δ*susD* (BT_3701) | 3701-5F | GCTGCAGGAATTCGATATCAggacttctccgcttgcagtcatggcaggag |
| | 3701-5R | caatttatcatgaaaacaaaatatatcaatgaaggctataaataaccaag |
| | 3701-3F | cttggttatttatagccttcattgatatattttgttttcatgataaattg |
| | 3701-3R | GTCGACGGTATCGATAAGCggctttcaatgaaaatctgaaatgggaaaaaac |
| *ptsH* | Up.ptsH-CmFRT-5v1 | GTCGAACCGCCAGGCTAGACTTTAGTTCCACAACACTAAACCTATAAGTT GGGGAAATACAGTGTAGGCTGGAGCTGCTTCGAAGTTCC |
| | Up-ptsH-CmFRT-5v2 | GGCTAGACTTTAGTTCCACAACACTAAACCTATAAGTTGGGGAAATAC AGTGTAGGCTGGAGCTGCTTCGAAG |
| *crr* | dn-crr-CmFRT-3v1* | CCGCTGGCGGAAGCATAAAAAAAATGGCGCCGATGGGCGCCATTTTTCA CTGCGGCAAGAACATATGAATATCCTCCTTAGTTCCTATTC |
| | dn-crr-CmFRT-3v2* | GAAGCATAAAAAAAATGGCGCCGATGGGCGCCATTTTTCACTGCGGCAA GAACATATGAATATCCTCCTTAGTTC |
| *ptsH* | Fw_verif_ptsH_crr | CCTTTTTAGGTGCTTTTTTGTGG |
| | Rv_verif_ptsH_crr* | CTCCGGTAATGAAGAAAATCAGG |
| *malE* | Fw_verif_malE | CAGACCTAGCACTGACC |
| | Rv_verif_malE* | CGAGATTGATATCTTTCGATACC |

**Table S3:** Model Parameters

| Parameter | Value | Description |
|---|---|---|
| $L$ | 1.0 | Normalized cavity length |
| $T_{\text{census}}$ | 20.0 | Total census time for AUC calculation |
| $r_{B,\max}^{\text{Gluc}}$ | 0.7 | Max growth rate of $B$ on Glucose |
| $r_{B,\max}^{\text{Am}}$ | 1.0 | Max growth rate of $B$ on Amylopectin |
| $r_{C,\max}$ | 1.3 | Max aerobic growth rate of $C$ |
| $f_{\text{ana}}$ | 0.2 | $C$ anaerobic growth fraction |
| $K_B^{\text{Gluc}}$ | 0.5 | Monod constant for Glucose ($B$) |
| $K_C^{\text{Gluc}}$ | 0.2 | Monod constant for Glucose ($C$) |
| $K_B^{\text{Am}}$ | 0.5 | Monod constant for Amylopectin ($B$) |
| $K_C^{\text{Ox}}$ | 0.5 | Monod constant for generic compound ($O$) ($C$) |
| $Da_{B,0}^{\text{Gluc}}$ | 0.7 | Baseline Damköhler for Glucose ($B$) |
| $Da_{C,0}^{\text{Gluc}}$ | 1.0 | Baseline Damköhler for Glucose ($C$) |
| $Da_{C,0}^{\text{Ox}}$ | 10.0 | Baseline Damköhler for compound ($O$) ($C$) |
| $Da_{B,0}^{\text{Am}}$ | 0.1 | Baseline Damköhler for Amylopectin ($B$) |
| $Pa_{B,0}^{\text{Gluc}}$ | 1.0 | Baseline production rate of Glucose |
| $G_0$ | 0.0 | Glucose concentration at $x = 0$ |
| $A_0$ | 1.0 | Amylopectin concentration at $x = 0$ |
| $\alpha$ | 1.0 | Compound ($O$) concentration at $x = L$ |
| $K_{\text{density}}$ | 5.0 | Baseline carrying capacity |
| $\varepsilon$ | 1.0 | Initial biomass seed density |
| $K_{\text{env}}$ | 0.1 | Compound ($O$) sensing half-saturation |
| $n$ | 4 | Hill coefficient for environmental modifier |
| $\lambda$ | 1.0 | Stoichiometric yield of Glucose from Amylopectin |
